# Walking Produces Oscillations in Orientation and Direction Perception at the Step Rate

**DOI:** 10.64898/2026.10.20.760552

**Authors:** Cameron K. Phan, Reuben Rideaux, David Alais

## Abstract

Walking is one of the most stereotyped oscillatory movement able-bodied individuals produce. It is highly consistent within and across individuals, with people taking ∼5000 steps a day on average at approximately 2 Hz. This consistency and its ubiquitous role in daily living make walking an ideal behaviour for investigating perception under active conditions. Across two virtual reality experiments, we tested whether stride-based oscillations, as seen in visual detection performance, also occur for visual orientation and motion direction discrimination. Participants followed a simulated screen along a linear path in virtual space while visual stimuli were presented on the screen. Group-level performance in both visual tasks oscillated at approximately two cycles per stride. Peak performance in visual orientation identification aligned with the late swing phase of each step, whereas peak performance in motion direction identification aligned with the late stance or early swing phase. These findings extend recent work showing that visual perception is modulated at the step rate during locomotion.

## Introduction

Walking is the most consistently observed and stereotyped active movement, with easily recognized gait phases that can be inferred from changes in physical indicators such as head height and centre-of-mass location (Hwang & Effenberg, 2021; Kharb et al., 2011; Rakovac, 2021). Head height varies approximately sinusoidally over the gait cycle, with peaks occurring in swing and troughs in stance (Hwang & Effenberg, 2021; Rakovac, 2021). Interaction between walking and vision has been investigated in the past, whether it be the influence of visual input on walking patterns (Prokop et al., 1997; Takamuku & Gomi, 2021; Warren et al., 2001) or the effect of walking on visual perception (Durgin et al., 2005; Pelah & Barlow, 1996; Yabe & Taga, 2008). However, the visual input manipulated or perceived in these studies were global scenes that would provide self-motion cues (optic flow), not individual moving stimuli. Additionally, these studies did not examine whether any changes in visual perception were linked to the gait cycle. Recent studies have shown that perceptual abilities exhibit sinusoidal modulation when measured during walking (Clouston et al., 2025; Davidson, 2023; Davidson et al., 2024; Phan et al., 2025). Detection performance for visual targets increases during swing and declines during stance (Davidson et al., 2024; Phan et al., 2025). These modulations are not restricted to perception: motor actions, such as the trigger pulls used to respond to visual or auditory stimuli, are entrained to footfalls, and manual tracking of an object is more precise during the swing (Davidson, 2023; Davidson et al., 2024; Phan et al., 2025). The existence of gait-related modulations in both motor and perceptual domains suggests that action-related modulation may be a prevalent feature of many perceptual abilities.

The current study employed a free-walking paradigm using wireless virtual reality (VR) technology and tested for gait-linked modulations of an orientation identification task and a motion identification task. The procedure was adapted from the flash-detection task used by Phan et al. (2025) which compared central and peripheral locations. Here we retained the visual location manipulation from Phan et al. to test whether the lead in stride-based modulation for central targets they observed would manifest differently between transient moving and static objects (relative to a simulated screen). The other gait effects noted above (perceptual sensitivity peaking in swing and motor responses more likely in stance) did not depend on eccentricity and showed no hemifield interactions (Phan et al., 2025). By substituting with oriented Gabor stimuli and measuring orientation judgments, our understanding of stride-based modulations of vision are expanded beyond contrast sensitivity or detection thresholds. Orientation, in particular, is of interest because it is essential for reaching and grasping, and because of its role as the predominant feature encoded in primary visual cortex. Furthermore, the addition of internal drift to the Gabor stimuli facilitates the study of visual object motion during walking. This form of local motion, while computationally resolvable in V1, evokes activity in, and may involve, the higher-level MT/V5 area (Albright & Stoner, 1995; Zaharia et al., 2019) and thus provides an opportunity to investigate the possibility of walking-induced modulations propagating through the visual hierarchy (Phan et al., 2025). Beyond its neural implications, motion is an object feature that is pivotal to safe and effective activity, particularly for navigating crowds and in sports (e.g., fielding in cricket and baseball).

How stride-based modulations occur in visual perception has been a point of discussion in the limited research to date (Davidson et al., 2024; Phan et al., 2025; Szekely & MacNeilage, 2025). One intuitive source for the variation in visual sensitivity is the modulation in the retinal image (Matthis et al., 2022; Szekely & MacNeilage, 2025). To maintain stable fixation, the moving observer’s eyes are required to rotate in opposition to the movement; this is known as the vestibular ocular reflex (VOR). For the orientation and motion direction tasks of the current study, retinal curl would increase difficulty and lower accuracy when the retinal image of the Gabor stimulus is rotated towards verticality (Matthis et al., 2022). This would occur when the head roll or sway are toward the same direction as the stimulus orientation, producing counterphase modulations between stimuli oriented congruently and those oriented incongruently to supporting foot and/or sway. Another VOR-induced change in retinal image is divergence (Matthis et al., 2022), in which the image is shifted further from the fovea (i.e., increased eccentricity) as the head sways towards the same side as the stimulus, relative to fixation. These VOR-induced changes in eccentricity would produce modulations in visual acuity and, consequently, counterphase accuracy modulations between stimuli presented in the visual field ipsilateral and contralateral to the supporting foot. Retinal image-based accounts of stride-based modulations in visual performance present predictions that are tested in the current study.

Another possible source of stride-based, or more general movement-based, modulations is that the movement itself, rather than the consequent retinal changes, induces oscillations in the visual cortex. This is supported by findings of synchronising oscillations in M1 and V1 following voluntary action (Benedetto et al., 2021). If stride-based modulations originate from the act of walking itself, they should not show the differences between lateralised stimulus features expected from retinal image changes (i.e., orientation alignment with head, lateral visual field). Instead, if the modulation begins in V1, a shift in phase between the orientation and motion direction task performance would indicate travelling wave behaviour (Fakche & Dugué, 2024; Muller et al., 2018; Phan et al., 2025; Sokoliuk & VanRullen, 2016), most likely from V1 to MT/V5.

Additionally, the motion direction identification task presents an opportunity to test the influence of self-generated movements on two known phenomena. First, motion perception exhibits biases that depend on observer-relative motion. For example, when the observer’s self-motion is opposite to an object’s motion, the object appears to move faster and be more salient (Fetsch et al., 2013; Hogendoorn et al., 2017). During the gait cycle, a walking individual’s head shifts up and down when entering and exiting the swing phase, respectively. Therefore, we expect greater likelihood of correct motion direction perception when the individual enters a phase in which their head moves opposite to the vertical direction of the internally drifting Gabor stimuli. The second phenomenon is the dichotomy between central and peripheral vision for static and motion stimuli, where peripheral vision is more sensitive to motion (Lewis et al., 2011). Any differences in perceptual modulation between centrally and peripherally presented stimuli would provide insight into the prioritization of visual feature processing across the retina during walking.

While higher frequency components are involved in the biomechanics of walking, such as the jolt in vertical height at heel-strike (Szekely & MacNeilage, 2025) and the dynamics of lower-limb joints (Tesio & Rota, 2019; D. A. Winter, 1984), stride-induced modulations in rotation and translation of the head are primarily low frequency (i.e., ∼1-2 Hz or ∼1-2 cycles per stride) (Grossman et al., 1988; Imai et al., 2001; Matthis et al., 2022; Szekely & MacNeilage, 2025; D. Winter, 1995). Likewise, walking observers’ gaze rotates and their pupils dilate at comparably low frequencies (Imai et al., 2001; Szekely & MacNeilage, 2025). Hence, in line with previous studies finding stride-based modulation of visual sensitivity (Davidson, 2023; Davidson et al., 2024; Phan et al., 2025), we expected modulations across the gait cycle to manifest at around two cycles per stride, which served as our frequency of interest.

## Methods

### Design

Experiment 1 employed a four-way 2 (Motion: Stationary vs Walking) × 2 (Visual Eccentricity: Central vs Peripheral) × 2 (Visual Field: Left vs Right) × 2 (Orientation: Left vs Right) repeated-measures design. An additional Drift Direction (Up vs Down) variable was implemented in Experiment 2, rendering it a five-way design. Factorial designs were employed to account for any potential confounds between independent variables. Comparisons of interest were between the levels of each independent variable.

Walking speed was set by the simulated display screen in the virtual environment positioned in front of the observer and centred at roughly eye level which traversed a 9.5 m distance in 9 secs (**Figure 1a, b**). Following the screen resulted in a walking speed of 1.056 m/s, slightly slower than typical human walking speed of approximately 1.4 m/s over long distances in natural environments (Hausdorff et al., 1996; Matthis et al., 2018). The slower walking speed was found to be a comfortable pace in our previous work, and appropriate given the uncertainty and unfamiliarity some participants may have experienced in a VR environment (Davidson et al., 2024). Target eccentricity was calculated for each presentation in real-time, based on the virtual viewing distance to the simulated screen. Central targets were displayed 3.7° (± 0.5°) to the left or right of fixation with no vertical offset, and peripheral targets were displayed 12° (± 0.5°) from fixation (**Figure 1e**). The eccentricity was calculated with the assumptions that participant gaze was perpendicular to the virtual display screen and that fixation was maintained. Prior to analysis, post-hoc corrective recalculation of target eccentricity was performed and targets were reassigned to two eccentricity ranges (See *Eye Movement Data*).

**Figure 1.**
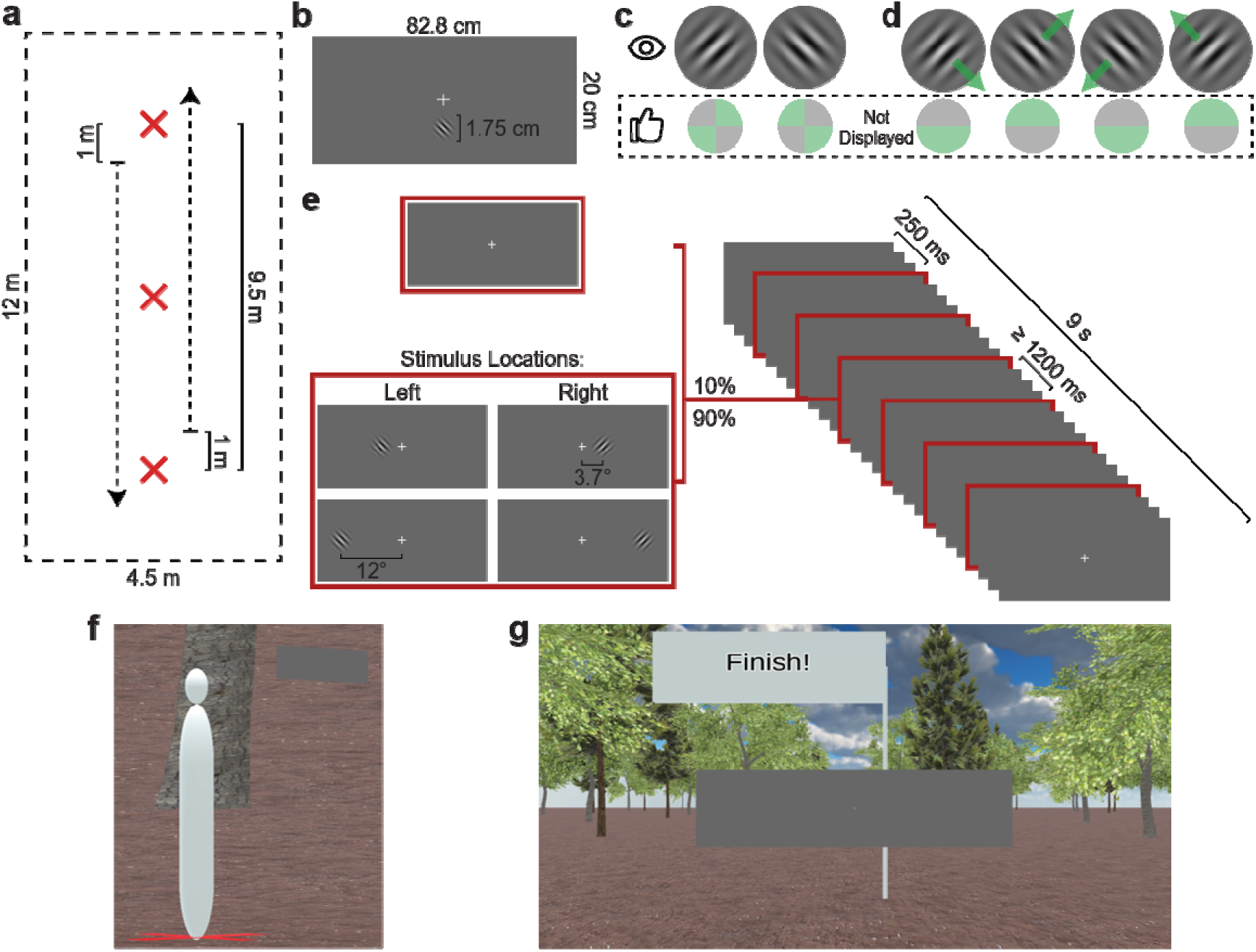
Schematic of trial sequence. (**a**) Simplified overhead layout of virtual walkway with the 9.5 m path transversed by the simulated display screen (arrows). The top and bottom red crosses indicate starting and ending positions for participants’ walking paths. The red cross in the centre indicated the participants’ position for stationary trials. (**b**) The dimensions of the simulated screen and target stimuli in layout of pre-sequence display. Participants pulled both left and right controller triggers to begin the trial sequence. (**c**) Full contrast examples of Gabor stimuli used as targets for the orientation judgment task of Experiment 1. Corresponding correct response quadrants on controller thumbpad shown below (not displayed to participant). (**d**) Full contrast examples of target stimuli for the motion direction judgment task of Experiment 2. Arrows indicate internal drift of the stimuli. Corresponding correct response hemifield of controller thumbpad shown below (not displayed to participant). (**e**) Example of 9 s trial sequence structure with 6 stimulus presentations (sequences could contain 5 or 6 presentations). 10% of stimulus presentations were omitted while 90% were equally distributed among the four stimulus location conditions (presented to the left). (**f**) Third-party perspective of simulated display screen relative to a stationary observer (1.625 m tall blank avatar; not simulated during experiment). (**g**) First person perspective of simulated observer in (**f**).

To investigate the interaction between lateral features of walking without the added complication of real-time determination of the stepping foot, we employed post-hoc labelling of targets. Targets presented on the same side of fixation as the supporting foot were labelled as ipsilateral and those on the opposite side were contralateral. Additionally, targets oriented towards the supporting foot (relative to verticality) were labelled as congruently oriented and those oriented to the swinging foot as incongruently oriented.

Unlike the simple yes–no responses used in prior studies (Davidson et al., 2024; Phan et al., 2025), responses on the circular trackpad of the controllers were mapped to quadrants (Experiment 1; **Figure 1c**) or vertical hemifields (Experiment 2; **Figure 1d**). Because of the bidirectionality of grating bars, the diagonal quadrant responses in Experiment 1 render the task a two-alternative response task. The limited response window and target contrast manipulation introduce opportunities for targets to not be detected at all in addition to assessing orientation and motion direction identification. A lack of response likely indicates a target that has not been detected. Incorrect responses indicate detection, but an incorrect perception of orientation (Experiment 1) or motion direction (Experiment 2).

### Participants

41 undergraduate psychology students participated in Experiment 1, while 33 participated in Experiment 2. Both samples were recruited from The University of Sydney, where each participant completed a scheduled one-hour experiment session for course credit, regardless of performance or completion of all experimental sequences. As per criteria below, 13 and 16 participants were excluded for Experiments 1 and 2, leaving final samples of *N* = 28 (*M_Age_* = 20.3 years, *SD_Age_* = 1.6, range = 18 – 23; 27 right hand dominant) and *N* = 17 (*M_Age_* = 19.8 years, *SD_Age_* = 2.8, range = 18 – 30; 15 right hand dominant), respectively. Each participant provided informed consent before commencing and had normal or corrected-to-normal vision. The study protocol was approved by the University of Sydney Human Research Ethics Committee (HREC 2021/048) and conducted in accordance with the Declaration of Helsinki.

### Apparatus and Materials

An HTC Vive Pro Eye head-mounted display (HMD) was used to display the virtual environment and two wireless HTC Vive Controllers (2018) were used for collecting responses. The HMD contained dual 3.5-inch high-resolution OLED displays (1440 x 1600 pixel resolution, 90 Hz refresh rate) with 110-degree field of view. Positions of the HMD and controllers were tracked in three-dimensional space at 90 Hz resolution, using five HTC Vive Base Station 2.0’s enclosing an open 4.5 m x 12 m rectangular space (**Figure 1a**). To allow the utilization of such a large space and unencumbered walking conditions, the virtual reality environment was transmitted to the HMD using the HTC Vive Wireless Adapter for Vive Pro. The SRanipal Runtime (Version 1.3.2.0; SDK 1.3.6.8) was used to handle eye tracking data collection by the integrated Tobii eye-tracker within the HMD, sampled at 90 Hz.

The virtual reality (VR) environment and experimental procedure were designed and presented within Unity (Version 2020.3.14f1), using the SteamVR Plugin (Version 2.7.3; SDK 1.14.15) on a Windows 11 PC (2 × 16 GB, DDR5 4400 MHz) with a 12th Gen Intel Core i7-12700K processor (3.60 GHz), and an NVIDIA GeForce RTX 3070 (8 GB, GDDR6) graphics card. The VR environment consisted of a clear space, that matched the dimensions of the tracked physical space, set in an open outdoor scene sparsely populated with trees with a single simulated natural light source. The trees, ground texture, and skybox used to create the outdoor environment were all free assets available on the Unity Asset store.

### Stimuli

All task-relevant visual stimuli were presented on a grey (RGBA 102, 102, 102, 255; 0.4, 0.4, 0.4, 1.0) rectangular screen with simulated dimensions of 20 cm × 82.8 cm (45° at 1 m horizontally; **Figure 1b,f,g**). This screen was smoothly transversing a linear path in real world space (not fixed to the head; **Figure 1a**). The target stimulus was a circular Gabor pattern (*frequency_grating_* = 1.92 cycles/deg, *SD_gaussian_* = 0.52°), simulated to be 1.75 cm (2.78° at 1 m), orientated 45° (± 5°) clockwise (right orientation) or anticlockwise from vertical (left orientation) (**Figure 1c**). For Experiment 2, the grating of the target stimuli was made to drift internally at 3 cycles per second, either (obliquely) upwards or downwards (**Figure 1d**).

To allow the largest range for oscillation in performance, a mean accuracy of 75% is optimal for two-alternative forced choice tasks, like those in the current study. Out of the features shared between the stationary and internally drifting Gabor targets (i.e., spatial frequency, orientation, contrast), contrast was the least problematic to adjust for task difficulty while retaining the ability to compare between the two perceptual tasks and the ability to disambiguate from previously found contrast sensitivity effects (Davidson et al., 2024; Phan et al., 2025). The reliance on capacitive touch detection of the controllers’ trackpads along with the relatively large surface area and capacitance of individuals’ thumbs precluded finer-grained estimation responses, which would be required if task difficulty were controlled using task features (i.e., orientation and motion direction). Alternatively, discrimination tasks involving two stimuli could have been employed, but the increased response time for such tasks would render the experiments impractical (e.g., participation would require several sessions, response times would span more steps, etc.) and introduce additional factors (e.g., relative location of the two stimuli, rendering two dynamic stimuli in wireless VR, etc.). Lowering spatial frequency would be analogous to slowing the internal drifting speed of targets in Experiment 2, which would require lower presentation times and larger temporal smearing along the stride cycle (Del Viva & Morrone, 1998), while the coarse orientation categorisation in Experiment 1 would require low spatial frequencies (high frequencies eliminated due to display resolution) equivalent to adjusting visibility (orientation would be converyed only at the edges of the aperture) (Burr & Wijesundra, 1991). If contrast-driven visibility is responsible for oscillations in performance, similar oscillations would be seen in response rate and accuracy (see *Design*). Hence, contrast was the most suitable feature to adjust for task difficulty.

The QUEST (Watson & Pelli, 1983) adaptive staircase algorithm was implemented to adjust the overall contrast of the Gabor patterns to produce an overall accuracy rate of 75% for each combination of motion and target eccentricity condition (i.e. unique staircases for central targets whilst stationary, central targets whilst walking, peripheral targets whilst stationary, and peripheral targets whilst walking). The initial slope (β) was set to 3.5, chance rate (γ) to.5, and lapse rate (δ) to.01 for each staircase, which had resolutions of.01 (i.e. 1%) for the range.01 to 1. As the QUEST procedure stabilized at 75% accuracy after ∼40 targets (∼ 8-10 walking sequences), a small jitter was added to each trial’s target contrast to prevent the staircases being resistant to variability in performance, particularly lapse rates, that would be expected for extended, not highly engaging tasks (Manning et al., 2018).

The virtual display screen would smoothly traverse linearly along the walking path at a constant velocity and constant height (calibrated to be at standing head height). This linear movement was independent of the participants’ head positions and was mapped as a straight path in world space coordinates (**Figure 1a**). On this fronto-parallel screen, a white (RGBA 255, 255, 255, 255; 1.0, 1.0, 1.0, 1.0) fixation cross was situated in the direct centre, and participants were instructed to maintain their gaze on it. Post-hoc analysis of eye tracking data allowed for verification of gaze fixation, both target and participant level exclusions, and recategorisation of target eccentricity (See *Eye Movement Data*).

### Procedure

After providing informed consent and given opportunity to ask questions, participants were introduced to the wireless VR apparatus, hand-controllers, power bank, the tracked physical space, and initial eye tracking calibration procedure before being fitted with the apparatus and given the hand-controllers.

Once the VR program was initiated, a 4-point calibration process for the HMD eye tracking was completed. Additional calibration was performed after any potential events that may have affected accuracy of tracking (e.g., signal drop-outs, breaks). Participants were then told to situate themselves in front of the virtual screen, standing on a virtual red cross 1 m away at the centre of the virtual environment (**Figure 1a**), and to read the task instructions The instructions outlined the requirement to always fixate on the fixation cross and to make speeded responses, pressing the quadrants on either hand controller’s trackpad corresponding to either end of the Gabor grating bands (**Figure 1c**). For Experiment 2, the task was to identify the vertical direction of the internal drift of the grating bands by pressing the corresponding hemifield of the trackpads (**Figure 1d**). Participants were also informed that to initiate each sequence, indicated by an example of the target at 100% contrast directly below the fixation cross, simultaneous trigger pulls on both hand controllers were required (**Figure 1b**). They were then asked to describe the task they were to complete to verify their understanding. If their understanding was correct, the participant was instructed to begin the first of three 9 s sequences while standing still. Otherwise, the participant was assisted in their understanding by alternate verbal instructions until they understood the task correctly and were given the opportunity to repeat the practice trial sequences. During this practice the targets presented were easy to detect, with a fixed supra-threshold constant in overall contrast (100%) and were not used to update any of the adaptive staircases.

Before beginning their first walking sequences, participants were instructed to stand on a red cross that indicated the starting location of the walking sequence (**Figure 1a**). The task was identical whilst walking, maintaining fixation with the additional instruction to maintain approximately 1 m between themselves and the display. For the first two sequences, the participant was accompanied to ensure their safety, and to verify this approximate distance was maintained comfortably whilst walking. Upon traversing the walking path, the participant was required to turn 180° before starting the following sequence, such that they return on the path they had walked to reach the previous start point. Once comfortable with the procedure of walking blocks, the participant was allowed to complete the remainder of the block independently, initiating the sequences themselves with dual-hand trigger pulls.

Participants completed 10 blocks, five stationary and five walking blocks, with the order of the blocks randomized except for the first block. Each block had twenty 9 s sequences, each of which had 5 or 6 targets presented, each target had a 10% chance of being withheld from presentation (**Figure 1e**). Targets were presented in one of four possible locations, either left or right of fixation and either centrally or peripherally, restricted to the virtual display screen. Targets were presented for 44 ms and were spaced by variable intertrial intervals with a minimum of 1210 ms. Instead, for Experiment 2, stimuli were presented for 198 ms with minimum intertrial intervals of 1210 ms. Targets were never presented during the first 253 ms of sequences. During sequences in walking blocks, the virtual display screen would smoothly traverse linearly along the walking path at a constant velocity and constant height, beginning motion when the participant started the sequence. After each walking sequence, if the participant was visibly further away or closer to the simulated screen than ∼1 m, they were told to walk faster or slower for the next walking sequence, respectively.

### Data Analyses

Each sequence provided 3D time-series data (x, y, z) for head position, head orientation, target position, gaze origin, and gaze direction. Individual steps were extracted based on head height, which follows a roughly sinusoidal pattern during walking (**Figure S1a,d**). Peaks and troughs in vertical head position correspond to the approximate swing and stance phases of the stride cycle, which we identified using a peak detection algorithm. Target performance data (i.e., hits, misses, false alarms and response times) were mapped according to target onset relative to nearest stride cycle percentile. Each stride cycle began with a trough in head height (presumably mid-stance) and included two steps, completing the cycle at the second trough at head height after the initial. Repeated-measures ANOVAs (see Supplementary Materials) were performed on the average of performance measures taken during walking across the conditions of the factorial design to inspect for systematic effects that may possibly drive modulatory differences, no effects of interest were found. All analyses were performed using custom Python (version 3.11.1) code, and ANOVAs were performed in JASP (version 0.18.0.0).

### Eye Movement Data

As Matthis et al. (2022) demonstrated and simulated, fixating during walking produces retinal curl and divergence due to the orbital rotation of the eye caused by the vestibular ocular reflex (VOR). To observe the potential contribution of these effects on performance, we decomposed the 3D gaze direction vectors into elevation and azimuth angles, analogous to pitch (**Figure S1b,e**) and yaw (**Figure S2a,b**), respectively. However, as convergent gaze measures were recorded, the interpretation of these angular changes needs caution as they are not measures of individual eye dynamics. When abnormal values occurred (i.e., > 1) in any of the three components of the normalised gaze direction vector, the observer was assumed to be blinking. Any target where the participant blinked during its presentation was excluded (Experiment 1: *M* = 0.14 targets, *SD* = 0.52, range 0 – 2; Experiment 2: *M* = 0.35 targets, *SD* = 1.06, range 0 – 4). Saccades were identified as any point when the 3D angular speed exceeded 35°/s and angular acceleration exceeded 9500°/s^2^. To avoid assigning small deviations as saccades, those smaller than 1° in magnitude were not counted as saccades. Saccades occurred during ∼19.3% (*M* = 77.18 targets, *SD* = 36.44, range 1 – 155) and ∼74.9% (*M* = 287.09 targets, *SD* = 47.36, range 183 – 340) of target presentations in Experiments 1 and 2, respectively. These gaze dynamics were analysed in the same manner as the head movements across the stride cycle (see *Gait Extraction*; **Figure S3**).

Using the projection of gaze (i.e., gaze direction from gaze origin), the eccentricity of the targets relative to fixation at stimulus onset was also calculated. For analysis, targets that were presented between 1.81° and 5.59° were assigned as central targets while those between 10.11° and 13.89° were assigned as peripheral. These ranges account for the jitter (± 0.5°) in presented eccentricities and intended radii of presented targets (± 1.39°). Any targets outside these ranges were excluded for central and peripheral target stride analyses (Experiment 1: *M* = 215.20 targets, *SD* = 12.72, range 185 – 235; Experiment 2: *M* = 168.65 targets, *SD* = 70.69, range 33 – 284). In the interest of maximising data density, targets determined to fall between the central and peripheral post-hoc categorization (i.e., 5.59° to 10.11° eccentricity) were retained for other target condition stride analyses (Experiment 1: *M* = 84.20 targets, *SD* = 37.84, range 16 – 176; Experiment 2: *M* = 90.03 targets, *SD* = 40.96, range 30 – 182).

### Gait Extraction

Stride-onsets were extracted based on the time-series of the vertical position (heave) of head-movement data as described in Davidson et al. (2024) and Phan et al. (2025). As walking shifts the centre of mass sinusoidally, troughs on the vertical axis of motion correspond to the double support stance phase of the gait cycle. Therefore, demarcated strides began at one trough in head height and ended two troughs later, encompassing two steps. Stride lengths were normalised for analysis by resampling the time-series data to 101 points (taken as percentage of progression: 0% – 100% inclusive). As such, stride progression reflects the execution of two consecutive steps. Using these demarcations, forward position (surge; **Figure S1c,f**), lateral position (sway; **Figure S2b,e**), roll (**Figure S2c,f**), yaw (**Figure S2a,b**), and pitch (**Figure S1b,e**) of the head were extracted and similarly resampled. To identify the frequency components of these dynamics across the stride cycle, they were analysed in the same manner as the perceptual performance measures (see *Performance Relative to Stride*), however without the additional binning beyond percentiles, due to the orders of magnitude more data these measures have (each frame vs discrete targets).

### Performance Relative to Stride

All target onsets were allocated to the percentile (from 1-100%) of the stride they occurred in. For stride analyses, performance was averaged within 20 linearly spaced bins, with zero overlap. Performance metrics analysed were response rate, proportion of responses made to presented targets, accuracy, proportion of correct orientation identifications out of total responses to presented targets, response time relative to target onset within progression (i.e., percentage), and response likelihood within progression.

Significant modulations were tested for each dependent variable for stride at the group level. Stepping from.1 to 10 cycles per stride, in increments of.1, raised cosine functions were fitted with the equation:

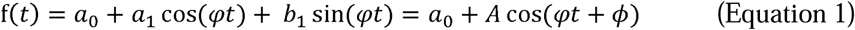

This is equivalent to fitting a minimal single-component (n = 1) Fourier transform where *φ* is the periodicity (cycles per stride), *t* is the progression of the stride (as a percentage), *a*_1_ and *b*_1_ are the coefficients of the cosine and sine components and *a*_0_ is a constant, the central value of the modulation (Ho et al., 2017; H. Zhang et al., 2019). The resulting raised cosine fits had amplitudes of A and shifts in phase of *ϕ*. This stepwise process is less prone than the fast Fourier transform (FFT) to spurious high-frequency components arising from discontinuities (i.e., mismatches between the start and end bin values), which are expected in behavioural data. For each forced fit at each frequency, the goodness of fit (*R*^2^) was used as an indication of strength. A resampling procedure was performed, where each of the 100 percentile bins were reallocated the measure of another bin, with replacement. These were then binned into their larger width bins and fits were performed as it was for the original data. This resampling was performed 1000 times with each resample having been fitted for the same frequencies as the original dataset. The *R^2^* values for the 1000 resamples at each fitted frequency were taken to represent the null distribution for strength of the modulations detected in the original data at the specific frequency, as the temporal intercorrelation of bins was effectively destroyed. The *R^2^* values for each frequency retained earlier were compared to the 95^th^ percentile of their respective null distributions, with those greater than this critical value taken to indicate a significant modulation.

### Exclusion Criteria

Three participants from Experiment 1 with data from less than four walking and less than four stationary blocks were excluded for insufficient data collection, caused by frequent breaks (*n* = 3). Head position data was visually inspected for aberrant sequences and abnormal gaits that may have occurred due to hardware malfunction, signal drop-out, or aberrant walking patterns (e.g., limping). Average of 0.37 sequences (*SD* = 0.66, range 0 – 3) and an average of 11.53 steps (*SD* = 11.46, range 0 – 28) were excluded per participant using this procedure for Experiment 1. For Experiment 2, an average of 1.31 sequences (*SD* = 0.60, range 1 - 3) and 9.79 steps (*SD* = 5.27, range 9 – 27) were excluded. One participant from each experiment no longer met the criterion for sufficient data (360 walking targets and 360 standing targets) and were excluded. Participants determined to not have fixated (i.e., kept gaze within 1°), according to the data analysis process, for at least 75% of their collective data points (Experiment 1: *n* = 3; Experiment 2: *n* = 3), were excluded. As mean performance approaches chance-level (i.e., 50%), the room for oscillatory variability reduces to 0, counter to the purpose of the current study. Hence, those with overall accuracy for each adaptive staircase, measured as the proportion of correct responses to total targets presented according to the corresponding staircase, less than 60% (Experiment 1: *n* = 6; Experiment 2: *n* = 12) were excluded. Similarly, those exceeding 90% accuracy would have been excluded, however, no participant achieved this. The high exclusion rate for low accuracy is most likely attributable to the combination of a lack of performance-based penalty, time-dependent reward for participation, and general simplistic psychophysical tasks (McKee, 1993; Verstraten, 2026). This is supported by the fact that there was heterogeneity in participants; majority were able to achieve mean performance within the inclusion range (i.e., 60 to 90%), some were incapable of completing all experimental sequences in the same session length (i.e., one hour), some completed all sequences but did not perform within the inclusion range. The first of these groups provide validation of the methods employed while the latter two highlight the need for future studies to adjust for variability in potential participant engagement. Further target-specific exclusions were applied based on eye tracking data (see *Eye Movement Data*).

## Experiment 1

### Results

Our wireless VR environment enabled an analysis of performance changes relative to stride progression by binning target onsets according to their stride-cycle phase (see Methods). This binning enabled the fitting of cosine functions and comparison to permuted data. We assessed the statistical significance of group-level modulations using a two-step permutation procedure, which allowed us to estimate the fit strength at chance-level (*R*^2^; see Methods). All modulations described below had raised cosine fits with *R*^2^ values exceeding the 95^th^ percentile of chance-level *R*^2^ values and were therefore deemed statistically significant, consistent with previous work (Ho et al., 2017; H. Zhang et al., 2019). For Experiment 1, participants responded to static Gabor stimuli by pressing on the trackpad quadrant corresponding to either end of the stimulus grating bars, rendering two of four options correct for all targets.

Accuracy (i.e., proportion of correct responses to all target presentations) for the orientation identification task is plotted in **Figure 2**. For five of the six conditions, accuracy showed a ∼2 cycles per stride modulation at the group-level. The exception was gratings presented in the ipsilateral visual field to the supporting foot. These ∼2 cycles per stride modulations showed similar phasic shifts, with accuracy peaking at the end of swing and declining as swing began.

**Figure 2.**
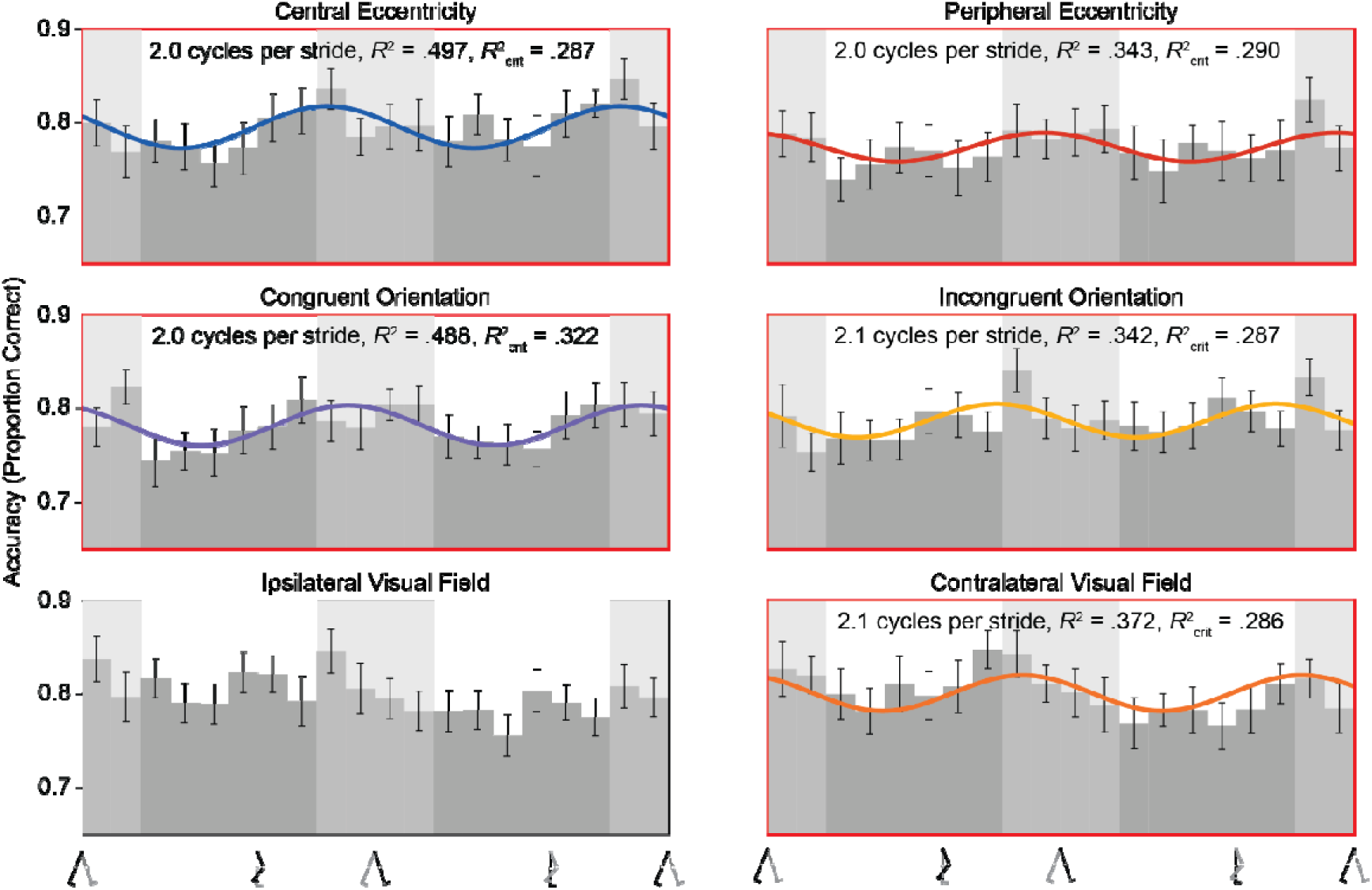
Accuracy for orientation identification across stride cycle. Group-level (*N* = 28) accuracy (proportion of correct responses out of total target presentations) data for the orientation identification task broken down by six target conditions; centrally presented, peripherally presented, congruently oriented to the supporting foot, incongruently oriented, presented in visual hemifield ipsilateral to the supporting foot, and presented in contralateral visual hemifield. Best fitting significant raised cosine fits are overlaid. Those significant fits in the frequency range of interest (1.5 – 2.5 cycles per stride) are bordered in red. Error bars indicate ±1 SEM. Light and dark grey columns indicate estimated stance and swing phases of the stride cycle, respectively.

**Figure 3** shows response rates (i.e., proportion of targets responded to) for the orientation identification task over the stride cycle. Response rates for the six target conditions oscillate idiosyncratically. The only ∼2 cycles per stride modulation arises for targets presented centrally and those tilted incongruently to the supporting foot (with participants more likely to respond in the stance phase; **Figure 3**). Importantly, response rate oscillations do not appear to be systematically related to the accuracy modulations in **Figure 2**, discounting the possibility that the accuracy modulations in **Figure 2** were driven by modulation in basic visibility.

**Figure 3.**
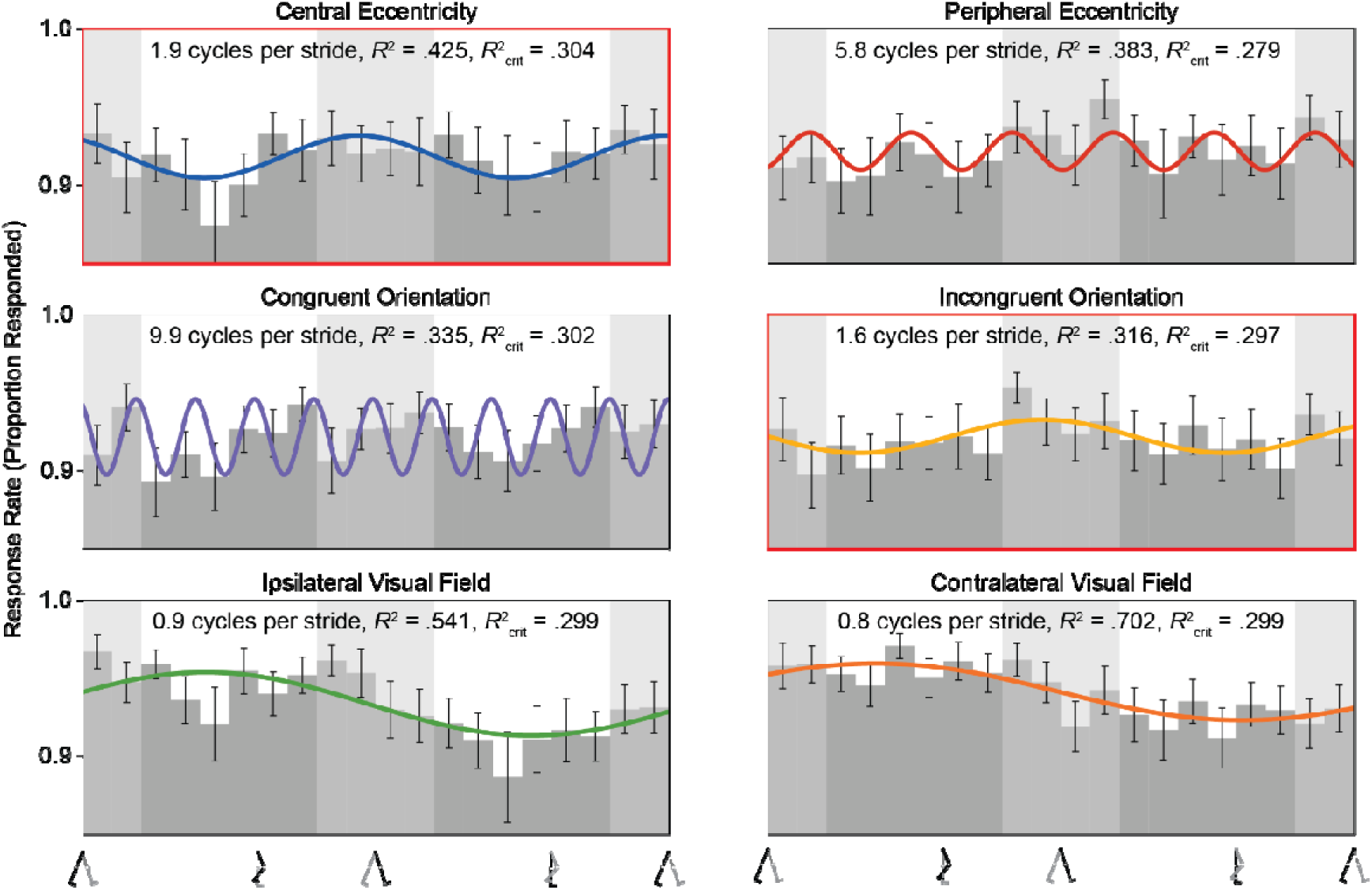
Response rate for orientation identification across stride cycle. Group-level (*N* = 28) response rate (proportion of target presentations responded to) data for the orientation identification task broken down by six target conditions; centrally presented, peripherally presented, congruently oriented to the supporting foot, incongruently oriented, presented in visual hemifield ipsilateral to the supporting foot, and presented in contralateral visual hemifield. Best fitting significant raised cosine fits are overlaid. Those significant fits in the frequency range of interest (1.5 – 2.5 cycles per stride) are bordered in red. Error bars indicate ±1 SEM. Light and dark grey columns indicate estimated stance and swing phases of the stride cycle, respectively.

Response times for the orientation identification task are shown in **Figure 4**. When targets were presented peripherally or were tilted towards the supporting foot, response times oscillated at 2.1 cycles per stride with the longest response times occurring in late swing or early stance phase. Given that the average duration of observers’ strides was ∼1.2 s (**Figure S4**), a single step lasted ∼600 ms, which places the responses to targets presented at the end of the swing phase near the end of the following step’s terminating stance phase. This is not in line with the peak in response likelihood seen for mid swing (**Figure S5**). There were significant response time oscillations for centrally presented targets, incongruently oriented targets, and those presented in the ipsilateral visual field to the supporting foot. However, these were at higher frequencies than those specific to the dynamics of walking (**Figures S1-S3**) and appear to be spurious.

**Figure 4.**
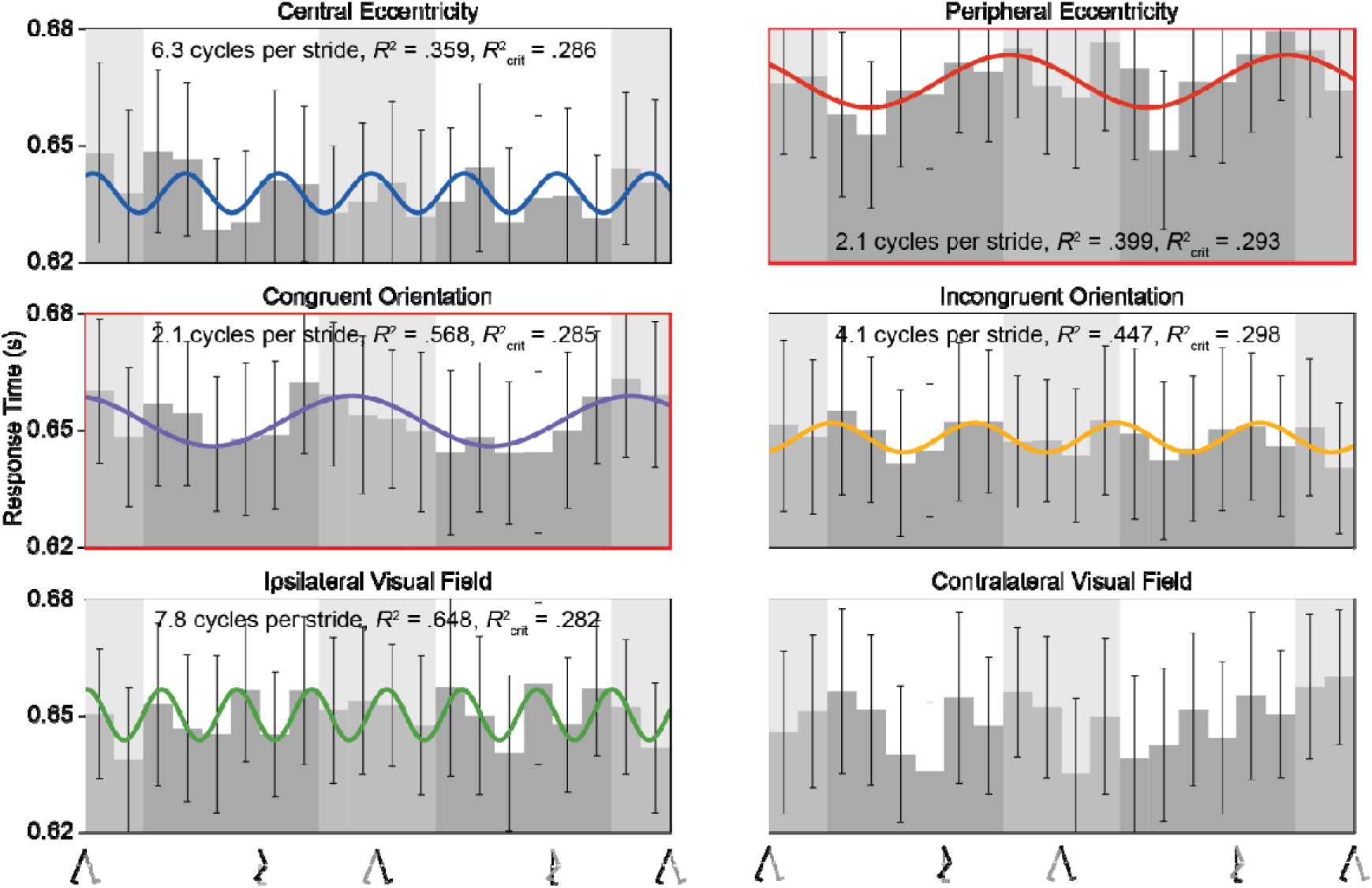
Response time for orientation identification across stride cycle. Group-level (*N* = 28) response time (s) data for the orientation identification task broken down by six target conditions; centrally presented, peripherally presented, congruently oriented to the supporting foot, incongruently oriented, presented in visual hemifield ipsilateral to the supporting foot, and presented in contralateral visual hemifield. Best fitting significant raised cosine fits are overlaid. Those significant fits in the frequency range of interest (1.5 – 2.5 cycles per stride) are bordered in red. Error bars indicate ±1 SEM. Light and dark grey columns indicate estimated stance and swing phases of the stride cycle, respectively.

### Discussion

Experiment 1 examined the effects of walking on visual orientation identification, focusing on performance changes within the stride cycle and these changes differed across visual locations and orientations. Participants made speeded button-press responses to indicate the orientation of the stimulus grating, effectively yielding a two-alternative forced-choice task.

Orientation identification accuracy was entrained to the rhythm of stride cycle for every combination of visual location and orientation, except for targets presented on the side of fixation ipsilateral to the supporting foot. Interestingly, we observed a modulation for stimuli presented contralateral to the supporting foot. Only one head transformation value, the roll of the head (**Figure S2c**), shows a similar or counterphase modulation to the one in performance for contralateral targets. However, this differential manifestation of stride-based modulation dependent on lateral presentation is not predicted by VOR retinal image accounts of head roll. As the head rolls towards one side, the eye should rotate in the counter direction, this would be seen as changes in the torsion of the eye and the polar angle location of target stimuli would be rotated about the fovea in the same direction of the head roll. Concerning retinal curl, targets oriented incongruently with the supporting foot will have retinal images oriented closer to verticality and those oriented congruently will be rotated towards the horizontal, which would produce counterphase modulations (not observed). However, these are speculations due to the difficulty in measuring torsion in the eye, which was not available in the VR apparatus and a limitation shared with Matthis et al. (2022). Since the location of the target was the feature of report and target orientations were quite away from either cardinal (i.e., vertical or horizontal), particularly in comparison to the small head roll modulation (< 1°), the VOR-induced changes in retinal image are not likely the source of the modulation for contralateral targets nor the lack of modulation for ipsilateral targets.

Phase alignment of all significant accuracy modulations was consistent - with accuracy peaking as individuals entered each stance phase of the stride - this may reflect global modulation of visual performance rather than a stimulus specific effect. Alternatively, the similarly aligned modulation of distance between observers’ gaze origin and the simulated screen may play a role in the accuracy modulations. However, the larger distance seen when accuracy is highest is counter-intuitive; being further away should decrease retinal image size and make targets more difficult to perceive, which was not reflected in response rates (a proxy for visual detection). Furthermore, this distance modulation was practically insignificant (< 0.4 mm; **Figure S7**) and its alignment with the accuracy modulation appears coincidental.

For the ∼2 cycles per stride modulation seen in the majority of conditions, the phase alignment was shifted in comparison to previous findings of stride-based modulations in visual detection (Davidson et al., 2024; Phan et al., 2025). The two tasks differ in that the visual detection task in those studies relied on simple contrast sensitivity whilst the orientation identification in Experiment 1 has the addition of orientation selectivity. The phasic shift between tasks may suggest that stride-based modulations of visual perception depend on task complexity. This raises the question of whether further increases in task complexity systematically shift stride-based modulations. Complexity could be added through additional stimulus feature combinations or the involvement of later visual cortices, such as making the task a global motion identification which requires the involvement of V5 (Albright & Stoner, 1995; Anstis et al., 1998; Newsome & Pare, 1988).

Similarly, there appears to be a shift in the response-making modulations observed in Experiment 1 in comparison with previous walking studies (Davidson et al., 2024; Phan et al., 2025). Instead of being more likely during the stance phase, response likelihood peaks during the swing phase. The source of this shift is most likely the greater decision making and motor production requirements a spatially mapped alternative-choice response has as opposed to a simple button press. Another difference is the selectivity of response-time modulation patterns. Due to greater response times in general for the orientation identification task, the previous-response time modulations found for contrast sensitivity in Davidson et al. (2024) and Phan et al. (2025) may not have been the major contributor to response time variability, resulting in only small modulations (∼10 ms) for two out of six target conditions.

## Experiment 2

### Results

In Experiment 2, participants responded to internally drifting Gabor stimuli by pressing on the vertical trackpad hemifield corresponding to the oblique vertical motion of the drift.

As shown in **Figure 5**, overall accuracy showed a ∼2 cycles per stride modulation at the group-level for half of the target conditions; peripherally presented, tilted congruently to supporting foot, contralaterally presented, and downwardly drifting targets. These ∼2 cycles per stride modulations show similar phasic patterns, with accuracy peaking at the start of the swing and declining as stance began. Response rates do not appear to explain these accuracy modulations: they only showed matching modulation for targets oriented to the supporting foot (**Figure 6**). Like in Experiment 1, the remaining target conditions exhibit higher frequency modulations that appear spurious.

**Figure 5.**
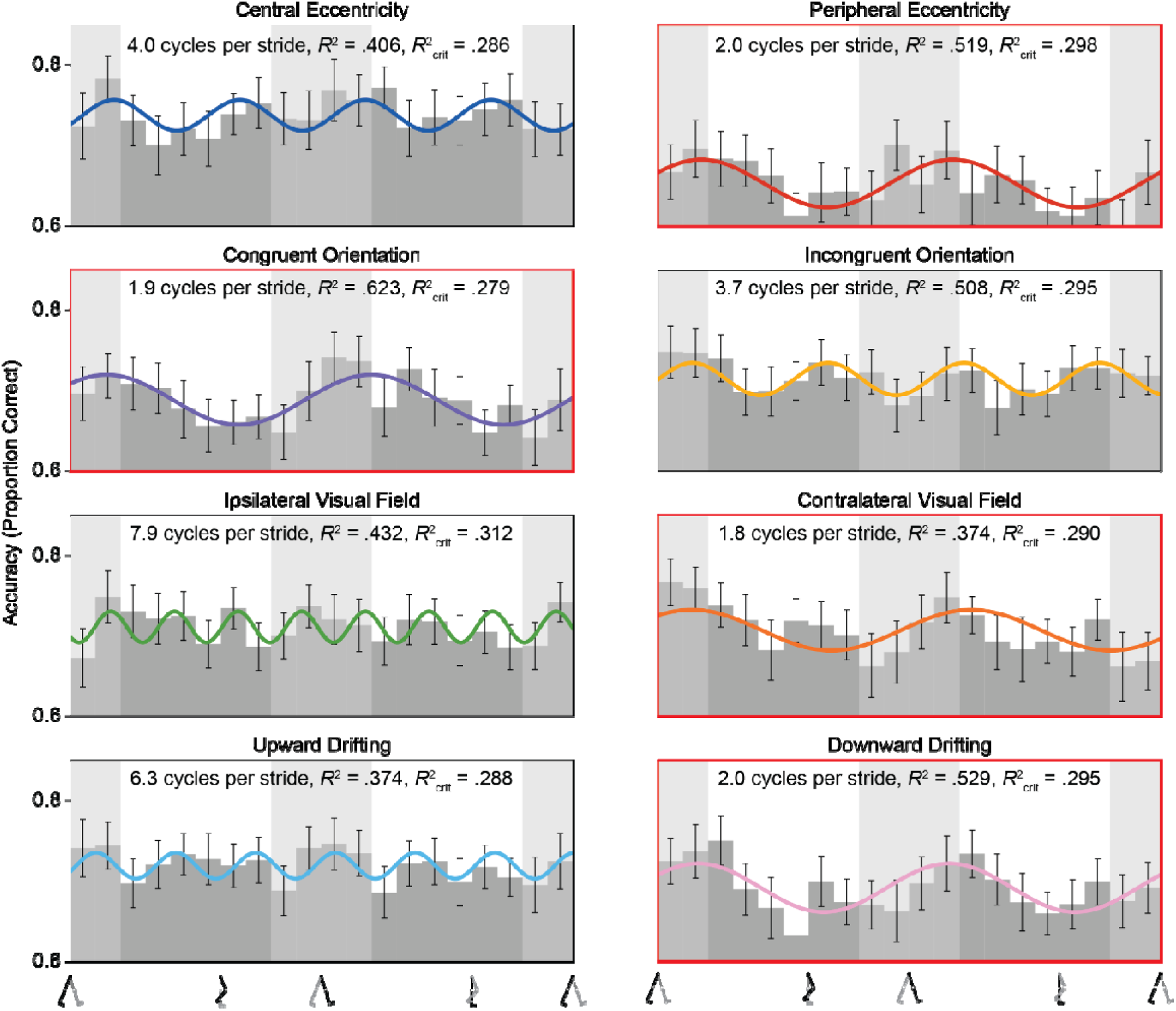
Accuracy for motion direction identification across stride cycle. Group-level (*N* = 17) accuracy (proportion of correct responses out of total target presentations) data for the motion direction identification task broken down by eight target conditions; centrally presented, peripherally presented, congruently oriented to the supporting foot, incongruently oriented, presented in visual hemifield ipsilateral to the supporting foot, presented in contralateral hemifield, upwardly drifting, and downward drifting. Best fitting significant raised cosine fits are overlaid. Those significant fits in the frequency range of interest (1.5 – 2.5 cycles per stride) are bordered in red. Error bars indicate ±1 SEM. Light and dark grey columns indicate estimated stance and swing phases of the stride cycle, respectively.

**Figure 6.**
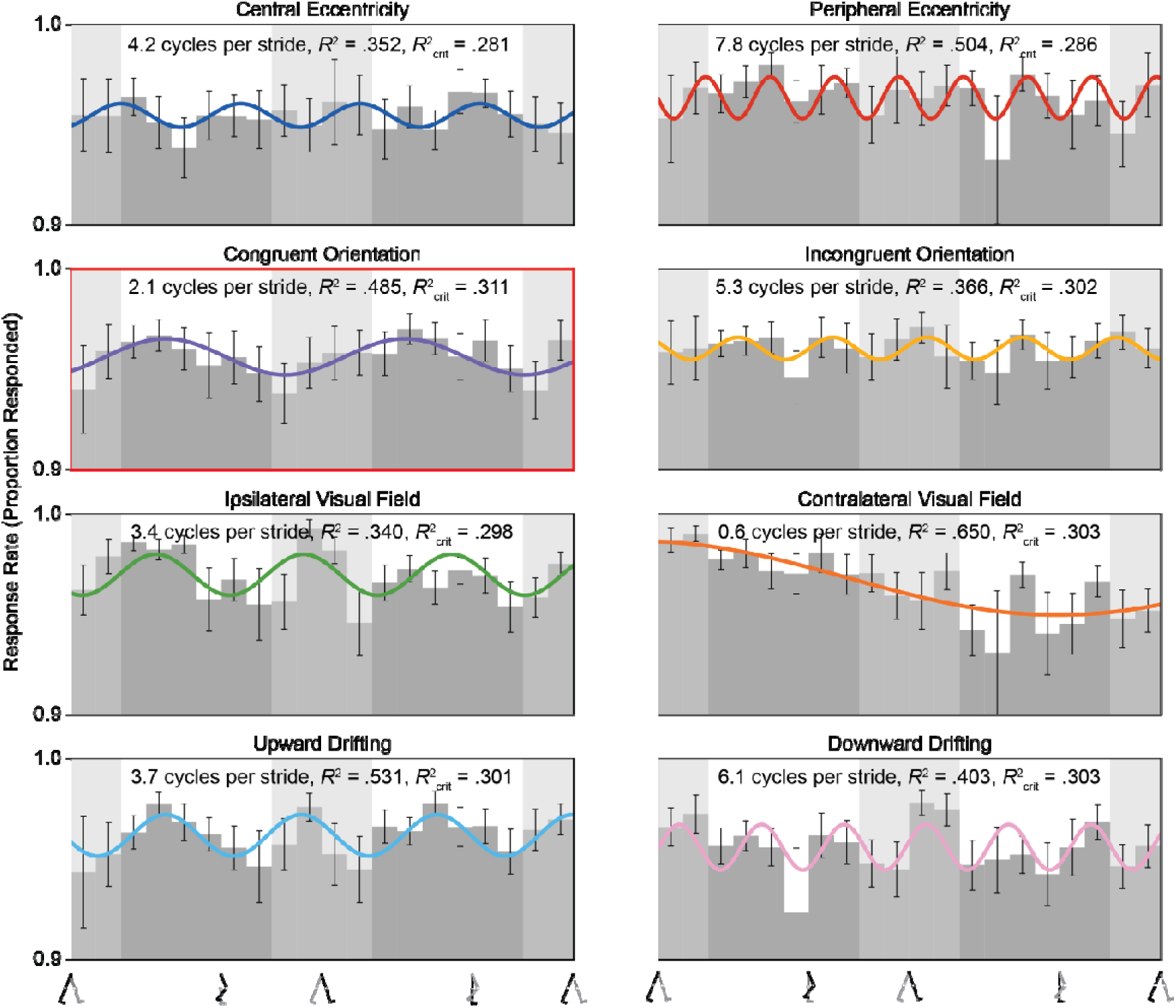
Response rate for motion direction identification across stride cycle. Group-level (*N* = 17) response rate (proportion of targets responded to out of the total target presentations) data for the motion direction identification task broken down by eight target conditions; centrally presented, peripherally presented, congruently oriented to the supporting foot, incongruently oriented, presented in visual hemifield ipsilateral to the supporting foot, presented in contralateral hemifield, upwardly drifting, and downward drifting. Best fitting significant raised cosine fits are overlaid. Those significant fits in the frequency range of interest (1.5 – 2.5 cycles per stride) are bordered in red. Error bars indicate ±1 SEM. Light and dark grey columns indicate estimated stance and swing phases of the stride cycle, respectively.

For the response time data (**Figure 7**), the significant modulations in the frequency range of interest were found for five out of eight target conditions. Except for contralaterally presented targets, these modulations show peak response times for targets presented in the late stance and early swing phase. These longer response times align well to produce responses during the mid swing phase as seen in response likelihood modulations (**Figure S6**). However, the shorter response times at the minima of the modulation do not produce the minima in response likelihood modulations. For contralaterally presented targets, response times peaked earlier, for targets presented in mid stance, and were misaligned with response likelihood modulations.

**Figure 7.**
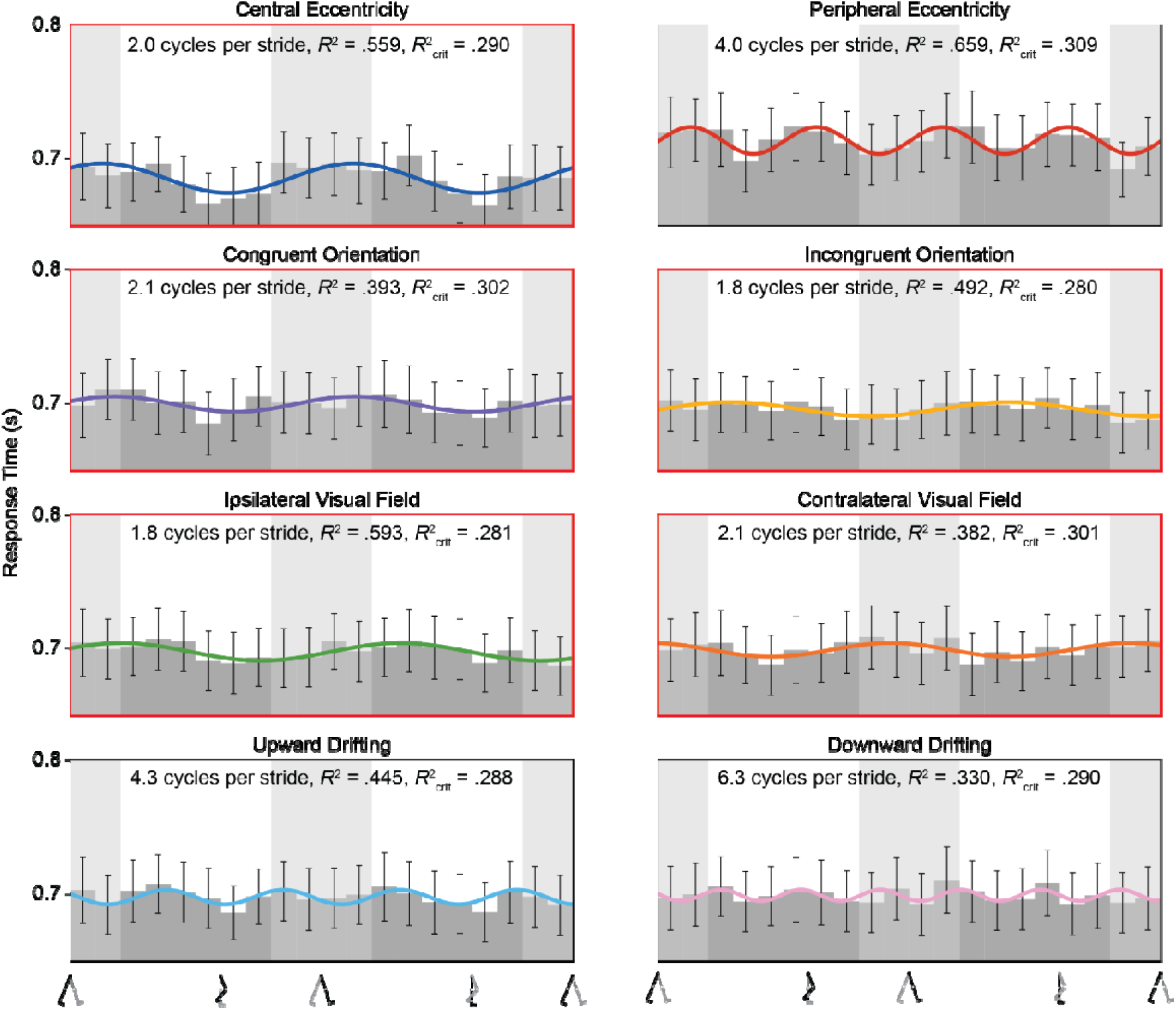
Response time for motion direction identification across stride cycle. Group-level (*N* = 17) response time (s) data for the motion direction identification task broken down by eight target conditions; centrally presented, peripherally presented, congruently oriented to the supporting foot, incongruently oriented, presented in visual hemifield ipsilateral to the supporting foot, presented in contralateral hemifield, upwardly drifting, and downward drifting. Best fitting significant raised cosine fits are overlaid. Those significant fits in the frequency range of interest (1.5 – 2.5 cycles per stride) are bordered in red. Error bars indicate ±1 SEM. Light and dark grey columns indicate estimated stance and swing phases of the stride cycle, respectively.

### Phasic Difference in Accuracy between Orientation and Motion Direction Identification

To investigate the differences in phasic shift (; Equation 1) of the accuracy modulations between Experiment 1 and 2, additional analyses were performed. Because we did not observe differences of interest in modulation features across target conditions in either experiment, we collapsed accuracy across all target conditions. Raised cosine fits were performed for each individual’s accuracy data. In a similar approach to the previous group-level analyses, goodness-of-fit *R*^2^ measures for shuffled resamples (*n* = 1000) were used to determine significance of these individual-level fits. Those who had significant two cycles per stride oscillations in each experiment were retained for comparison of phasic shift between tasks (orientation: *n* = 10; motion: *n* = 6; **Figure 8a**).

**Figure 8.**
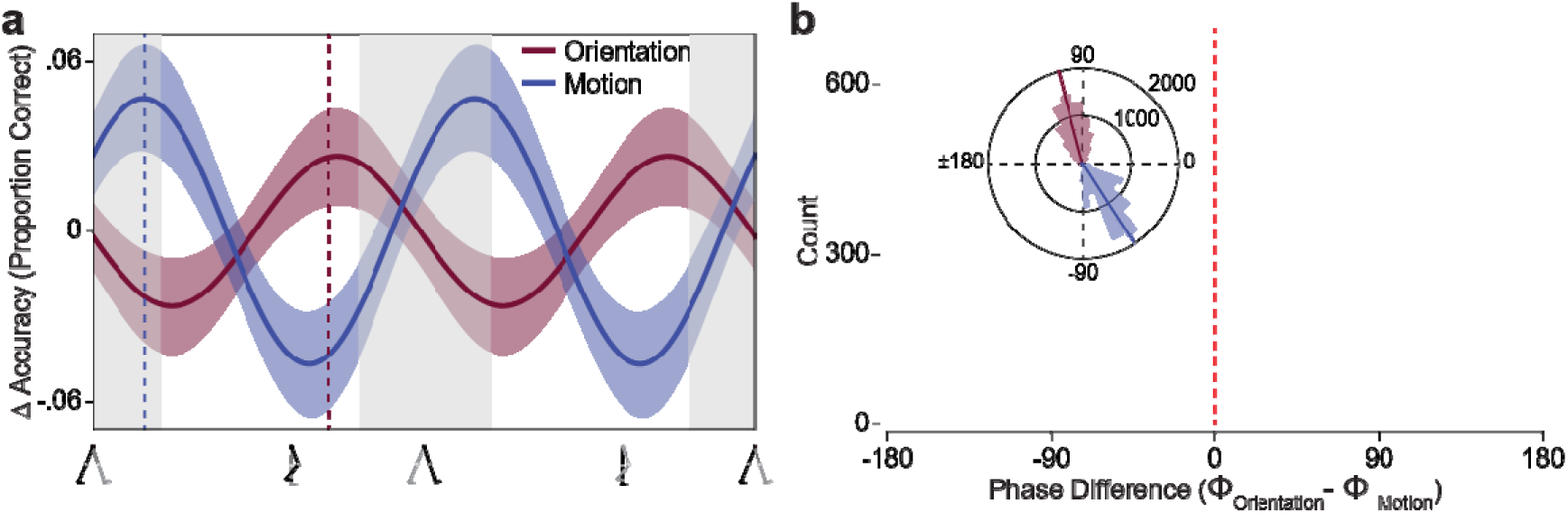
Accuracy modulations between the orientation and motion identification across stride-cycle. (**a**) Average raised cosine fit from participants with significant two cycles per stride oscillations in accuracy for the orientation identification task (*N* = 10) and motion direction identification task (*N* = 6). The orientation identification modulation is in a dark red and motion direction identification modulation in dark blue. Shaded regions indicate ± 1 SEM. Light grey columns indicate estimated stance of the stride cycle. Vertical dashed lines (in corresponding colours) indicate the approximate peak accuracy point along the first step. (**b**) Bootstrapped (*n* = 10,000) difference in phase means between the two tasks. Overlaid are the two bootstrapped phase distributions of each task.

To provide an approximation of the distribution of phase shifts for each experiment and the phase difference between tasks, a bootstrap resampling (n = 10,000) procedure was applied (**Figure 8b**). From this procedure, the phases of the two tasks’ accuracy modulations differed significantly (*ϕ_orientation_* — *ϕ_motion_* = 2.83 rad, 95% CI [0.92, 4.96]). Accuracy for the orientation identification task peaked at ∼71% (*ϕ_orientation_* = 1.81 rad, 95% CI [0.25, 3.64]) of the step cycle (late swing phase), whereas accuracy for the motion identification task peaked at ∼16% (*ϕ_motion_* =-1.01 rad, 95% CI [-2.45, 0.11]) of the step cycle (late stance phase).

### Discussion

Experiment 2 shifted the visual feature of interest from orientation to motion, increasing stimulus complexity and processing latency while retaining a two-alternative forced-choice response. Participants made speeded responses on a mapped thumbpad to indicate whether the grating’s motion direction was upwards or downwards.

Motion direction identification accuracy modulated at ∼2 cycles per stride for half of the conditions, one from each pairwise comparison. The remaining four conditions did not modulate at any frequency seen in the head or gaze dynamics (**Figure S1-3**) nor do their modulations appear to be nested within each step. In comparison to the accuracy modulations in orientation identification, the ∼2 cycles per stride modulations seen in Experiment 2 were shifted in phase, such that accuracy peaked as individuals left as opposed to when they entered the stance phase (i.e., modulation was later in Experiment 2 and near counterphase). These modulations align with that seen in head, but not gaze, pitch (**Figure S1e**), head roll (**Figure S2f**), and saccade probability (**Figure S3**). Like in Experiment 1, retinal image modulations as a result of head roll would be expected to produce counterphase modulations between lateral visual fields and orientation congruency, which was not observed. Instead, each of the pairings only had one condition with ∼2 cycles per stride modulations, a differentiation not supported by VOR accounts. Concerning head pitch, there is no real retinal image or vestibular account that would explain best performance when an observer’s head is pitched relatively upwards. This leaves one final candidate: saccades.

Perisaccadic suppression refers to the decreased visual sensitivity observed around the onset of a saccade, both shortly before and during the early initiation of eye movement. This would be expected to produce modulations in counterphase to those seen in Experiment 2, aligning better with the accuracy modulations seen in Experiment 1. Alternatively, the longer stimulus duration could have provided greater opportunity for reactive saccades to occur. This is unlikely as reactive saccades typically require ∼200 ms to initiate and would be expected to produce a peak in probability ∼17% in stride progression after the peak in accuracy (Robinson, 2022). Therefore, it appears the increased probability of saccades during target presentation in Experiment 2 is the result of a baseline rate of saccades across a longer stimulus duration. This is supported by the rate of target presentations with saccades being roughly equivalent in ratio to the stimulus durations of Experiments 1 and 2 (i.e., ∼4×).

As a feature dependent on a spatial element changing over the course of time, motion fundamentally requires minimum of two time point samples to be perceived, as opposed to the single time point needed for orientation (Del Viva & Morrone, 1998; Derrington et al., 2004). In V1, this feature tracking has been attributed to complex cell activity. Complex cells integrate activity of the orientation-selective simple cells and as a result of this hierarchical structure, have a greater latency (Hubel & Wiesel, 1962; Van Kleef et al., 2010). Beyond these early and small differences in latencies, there is additional difference in cortical areas activated in orientation and motion perception. The Middle Temporal (MT or V5) area demonstrates much greater activity when processing moving stimuli and weak activity for static stimuli (Raiguel et al., 1999). Two processes have been proposed as impetus for V5 activity, a parallel processing account where V5 receives output from subcortical areas (Holliday et al., 1997) and the traditional hierarchical model where input originates from V1 (Maunsell & Essen, 1983). The subcortical pathway to V5 has been associated with differences in response latencies between V1 and V5 below 100 ms (Foxe & Simpson, 2002; Lamme & Roelfsema, 2000; Raiguel et al., 1989). Difference in neural response latencies more appropriately matching the phasic difference of the accuracy modulations between Experiment 1 and 2 are those attributed to the pathway between V1 and V5, specifically feedback from V5 (Prieto et al., 2007). Therefore, if a cortical activity measure were to be incorporated, it would be probable that V5 would show a more delayed oscillation than V1 during walking.

As seen in Experiment 1, the response likelihood peaked during the swing phase. This repetition of phasic pattern is a strong replication but different to those seen for simple button-press responses in previous visual detection tasks. This further strengthens possibility of the driving factor for the phase shift being greater decision-making and motor production requirements for mapped alternative-choice responses. The majority of the ∼2 cycles per stride modulations in response time found for Experiment 2 showed longer response times at the end of the stance or early in the swing phase. However, these modulations could only account for the peak in response likelihood, not the low likelihoods in stance phases. As in Experiment 1, the longer response times required may have diminished the contribution of phase optimisation of response making to response time.

## General Discussion

From both experiments, the effect of stimulus orientation on visual abilities during walking can be observed. Regardless of whether orientation was directly relevant to the perceptual task (Experiment 1) or indirectly relevant (grating motion task in Experiment 2), alignment of the stimulus orientation to head orientation (i.e., roll; **Figure S2c,f**) did not play a role in modulation of task performance across the stride-cycle (as discussed for Experiment 1). Overall, both tasks produced accuracy modulations over the stride cycle. The phase difference between these modulations may reflect differences in processing requirements when determining a grating’s orientation and identifying its orthogonal motion when drifting. In both tasks, the accuracy modulations were similarly aligned even when dividing targets into different visual feature categories, suggesting a task-based modulation of percept across the stride cycle.

Focusing on the motion direction identification task in Experiment 2, the asymmetry in the influence of the stride cycle on accuracy between upward and downward drifting stimuli is unexpected. Only identification of the direction of downwardly drifting stimuli demonstrated stride-based modulation, peaking as individuals entered the swing phase, when their centre of mass and head height moved upwards. This greater accuracy could be due to greater, more detectable motion magnitude, explained by simple vector addition between the vestibular vector and the visual targets motion vector (Fetsch et al., 2013; Hogendoorn et al., 2017). Along the same line, it was expected that the accuracy would have peaked for upwardly drifting stimuli when the individuals centre of mass and head height moved downwards. However, absence of such an effect and its consequential stride-based modulation in accuracy for these stimuli weakens this account. Similarly, a response bias explanation is unlikely. If modulation in bias between responding upward or downward was responsible, we would expect counterphase modulations for upward and downward drifting targets. Even if both perception and response bias modulate separately, their summation would be the expected modulation in performance. Addition of modulations of the same frequency, two cycles per stride in this case, would simply result in a phase shifted modulation of the same frequency, not the non-significant higher frequency modulation seen for upward drifting targets. Furthermore, similar to orientation identification performance, consistent observation of similarly phase aligned modulations in other visual features suggests a non-selective stride-based effect for vertical motion direction.

Could the stride-based modulations in the current study be the result of retinal motion artefacts? Matthis et al. (2022) demonstrated that fixating on the ground plane during walking produces retinal curl and divergence due to orbital rotation of the eye caused by the vestibular ocular reflex. Both retinal curl and divergence increase with eccentricity from fixation and angular difference between the eye’s velocity vector and its gaze vector. This means curl and divergence are greater for ground plane fixation and much reduced for fixation on points well above the ground on the fronto-parallel plane. This was recently confirmed by Powell et al. (2026), who found that retinal curl was eliminated when optic flow cues were removed using blank walls, analogous to the fronto-parallel and roughly eye-level mid-grey screen that stimuli were presented on in the current study (**Figure 1f,g**). Even if we are to assume the highest retinal curl (∼ 0.2°) and divergence (∼ 0.9°) shown by Matthis et al. (2022), they would provide practically insignificant contributions to the 40° to 50° orientation and eccentricity (3.2° to 4.2° and 11.5° to 12.5°) of the target stimuli presented in the current study. Furthermore, by calculating actual target eccentricity on each trial and then excluding those that fall outside reasonable bounds, we effectively eliminate what little retinal divergence could have occurred. Retinal curl would exert a similarly small effect, adding a small component to Experiment 2’s Gabors’ drift rate of about 0.4% and would not materially alter thresholds (Lappin et al., 2009). It should be highlighted that these calculations are based on the values from 60° eccentricity of a single eye fixated on a textured ground plane, which provides rich optic flow cues. Considering the practically small variation we observed in head and gaze direction rotation (i.e., < 3°; **Figure S1b,e** and **Figure S2**) and that the lateral and forward translation in the head (the origin of the gaze) are insignificant (< 0.6 mm; **Figure S2b,e** and **Figure S7**), it can be assumed that the findings of the current study are not the result of VOR-driven changes on the retina.

Instead of retinal image changes, neuronal sensitivity may undergo modulations corresponding to those seen in the current study. Benedetto et al. (2021) found synchrony between oscillations in M1 and V1 after the execution of voluntary action using fMRI in human observers. These V1 oscillations were within the theta frequency band and were correlated with behavioural response accuracy. A key difference between walking in the current study and the keypress response in Benedetto et al.’s study is the automaticity and rhythmicity of the former (Clark, 2015), as opposed to the transient nature of the latter. Saccades are another, more automatic, action associated with changes in visual sensitivity. McFarland et al. (2015) found increased V1 activity but decreased stimulus selectivity in macaques shortly after their saccades. The activity had a temporal profile matching input from the lateral geniculate nucleus, which was consequently identified as the source of the selectivity decrement. If a similar mechanism were responsible for our findings, it would be expected to originate in areas associated with the automatic generation of walking movements, such as the brainstem or cerebellum (Morton & Bastian, 2004; Takakusaki et al., 2016) instead of the more preparatory M1 cortical area (Benedetto et al., 2021). Activity in the cerebellum can be non-invasively recorded using electrocerebellography (ECeG) (Todd et al., 2025) and has shown movement-related synchronisation of spectral power in the delta (0.5 to 4 Hz) band (Körmendi et al., 2021; Popovych et al., 2016), making it a highly suitable candidate for investigating stride-based modulations.

Both experiments conducted in the current study found novel and unique effects of stride cycle on response making. Instead of the previous response preference for the stance phase for simple visual detection (Davidson et al., 2024; Phan et al., 2025), both alternative-choice tasks showed response patterns that peaked during the swing phase. Furthermore, response times, previously found to peak during the stance phase, did not modulate to prioritise phase of response making. This most probably reflects a smaller contribution of response phase optimization in comparison to the longer decision making and response making times. A future study could verify this by employing a continuous tracking paradigm where the temporal delay associated with peak of response correlations could be taken to reflect stride cycle phase response optimisation (Ambrosi et al., 2022).

Considering previous stride-based modulations in visual detection alongside the current findings, there appears to be a phasic shift that depends on visual task complexity. Both visual detection and orientation identification occur in V1, the former is simple activation whilst the latter involves a percept-based response (Koch & Ullman, 1987; X. Zhang et al., 2012). Therefore, despite the same cortical area, the processing required is greater for orientation identification. Processing is greater again in the motion task as perception of visual motion can require involvement of V5, a later cortical area (Albright & Stoner, 1995; Anstis et al., 1998; Newsome & Pare, 1988). If stride-based modulations for performing these tasks are compared, phase is shifted in a manner consistent with the change in task complexity. Peak of performance is in the middle of the swing phase for visual detection, end of the swing phase for orientation identification, and beginning of the swing phase for vertical motion identification. Due to the cyclic nature, the beginning of the swing phase can be interpreted as either earlier or later. Hence, there is a need for the underlying cortical activity to be measured across the visual hierarchy to verify whether there is a stride-based modulation of visual cortical activity and, if so, whether it propagates along the hierarchy in a continuous macroscopic travelling wave manner (Muller et al., 2018).

Beyond probing further along the visual hierarchy, future studies could investigate orientation and motion perception in walking observers more directly. The trackpads used to collect observers’ responses in the current study rely on capacitive touch detection and are rather coarse in resolution, rendering analysis of estimation error inappropriate. Using alternative response devices or two-stimulus discrimination tasks, studies could measure modulation in precision or tuning across the stride cycle. Alternatively, other features of the Gabor stimuli employed could be the target of perceptual tasks (e.g., spatial frequency or speed of internal drift). There remains a myriad of features and perceptual tasks that could be investigated in walking, or moving, observers.

The current study has expanded previous findings of locomotion-driven modulations in visual perception. While the original stride-modulation paper by Davidson et al. (2024) used a simple visual detection task in central vision, here we include tasks involving greater complexity and targets presented in different visual field locations. Congruency of orientation, lateral visual field, and vertical direction of visual motion do not appear to undergo unique modulations during walking. However, it does appear that general perceptual task requirements affect the phase of stride-based modulations. Orientation identification, which simply involves V1, exhibited a peak in task performance earlier in the stride cycle than motion direction identification, which generates activity in the later cortical area V5. Together, the phasic and hierarchical differences suggest that stride-based visual modulation originates in lower cortical areas and propagates along the cortical hierarchy as a travelling wave. This could be tested in future research in a number of ways. One would be by employing neurophysiological measures such as electroencephalography that could reveal evidence of such propagation. Another approach would be to use higher-order stimuli such as faces and foods, both of which involve fusiform areas (Jain et al., 2023; Kanwisher & Yovel, 2006; Khosla et al., 2022), and more complex tasks such as evaluating face emotion or food categories.

## Supplementary Materials

### Average Performance in Orientation Identification While Walking

A four-way 2 (Visual Eccentricity: Central vs Peripheral) x 2 (Visual Field: Left vs Right) x 2 (Step Foot: Left vs Right) x 2 (Gabor Orientation: Left vs Right) repeated-measures ANOVA was performed on four dependent variables: response rate, accuracy, correct response time, and overall response time from target presentations during walking. For accuracy, there were no significant main effects nor any significant interactions, *p* >.05. For response rate, there was only a significant main effect of eccentricity such that participants were more likely to respond to targets presented centrally than peripherally, *F*(1, 27) = 10.315, η^2^ =.009, *p* =.003. For overall response time, there was a significant main effect of eccentricity with participants responding faster to central than peripheral targets, *F*(1, 27) = 23.500, η^2^ =.095, *p* <.001. Participants were also significantly faster to respond to targets presented in their right visual fields, *F*(1, 27) = 7.103, η^2^ =.021, *p* =.013. These main effects of eccentricity, *F*(1, 27) = 20.125, η^2^ =.081, *p* <.001, and visual field target location, *F*(1, 27) = 7.533, η^2^ =.022, *p* =.011, were significant for correct response times. There were no significant interactions of interest for response times.

### Average Performance in Motion Direction Identification While Walking

A five-way 2 (Visual Eccentricity: Central vs Peripheral) x 2 (Visual Field: Left vs Right) x 2 (Step Foot: Left vs Right) x 2 (Gabor Orientation: Left vs Right) x 2 (Drift Direction: Up vs Down) repeated-measures ANOVA was performed on the same four dependent variables as was performed on four dependent variables: response rate, accuracy, correct response time, and overall response time from target presentations during walking. Participants responded with greater accuracy when targets were presented centrally than when presented peripherally, *F*(1, 16) = 49.56, η^2^ =.062, *p* <.001. There were no other significant main effects on accuracy nor any significant interactions of interest. Participants were responded to a greater proportion of targets presented centrally than those presented peripherally, *F*(1, 16) = 4.80, η^2^ =.037, *p* =.044. There were no other significant main effects on response rate nor any significant interactions of interest. For overall response times, individuals were significantly faster to respond to targets presented centrally than those presented peripherally, *F*(1, 16) = 74.54, η^2^ =.153, *p* <.001. There was a significant interaction between the orientation of the target and the foot that initiated the step it was presented during, such that when the target was tilted to the same side as the foot response times were greater, *F*(1, 16) = 5.89, η^2^ =.005, *p* =.027. There were no other significant main effects on overall response times nor any significant interactions of interest. When excluding incorrect responses, response times retained the main effect of eccentricity, *F*(1, 16) = 79.05, η^2^ =.148, *p* <.001, and Step Foot x Gabor Orientation interaction, *F*(1, 16) = 10.03, η^2^ =.008, *p* =.006. There were no other significant main effects on the correct response times nor any significant interactions of interest.

---

**Figure S1.**
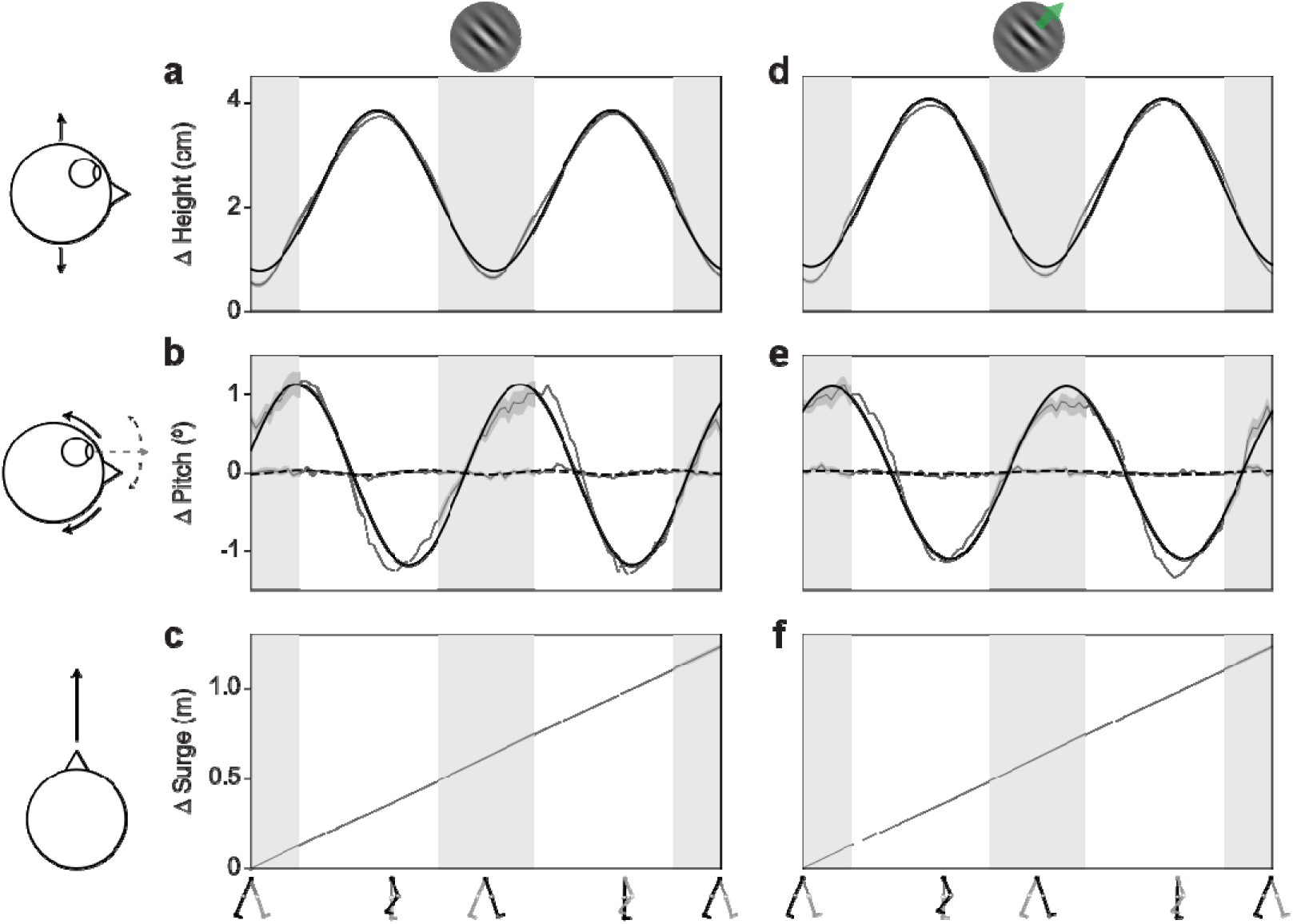
Height, pitch, surge across stride cycle. (**a**) Group-level (*N* = 28) change in head height across stride cycle for the orientation identification task. The stride cycle is comprised of two steps, beginning mid stance phase. (**b**) Same as (**a**), but for the change in pitch in head orientation (solid line) and gaze direction (dashed line). (**c**) Same as (**a**), but for the surge in head height. (**d**-**f**) Same as (**a-c**), but for the group-level (*N* = 17) data from the motion direction identification task. Best fitting raised cosine functions are overlaid for (**a, b, d, e**). Shaded regions indicate ±1 SEM. Light and dark grey columns indicate estimated stance and swing phases of the stride cycle, respectively.

**Figure S2.**
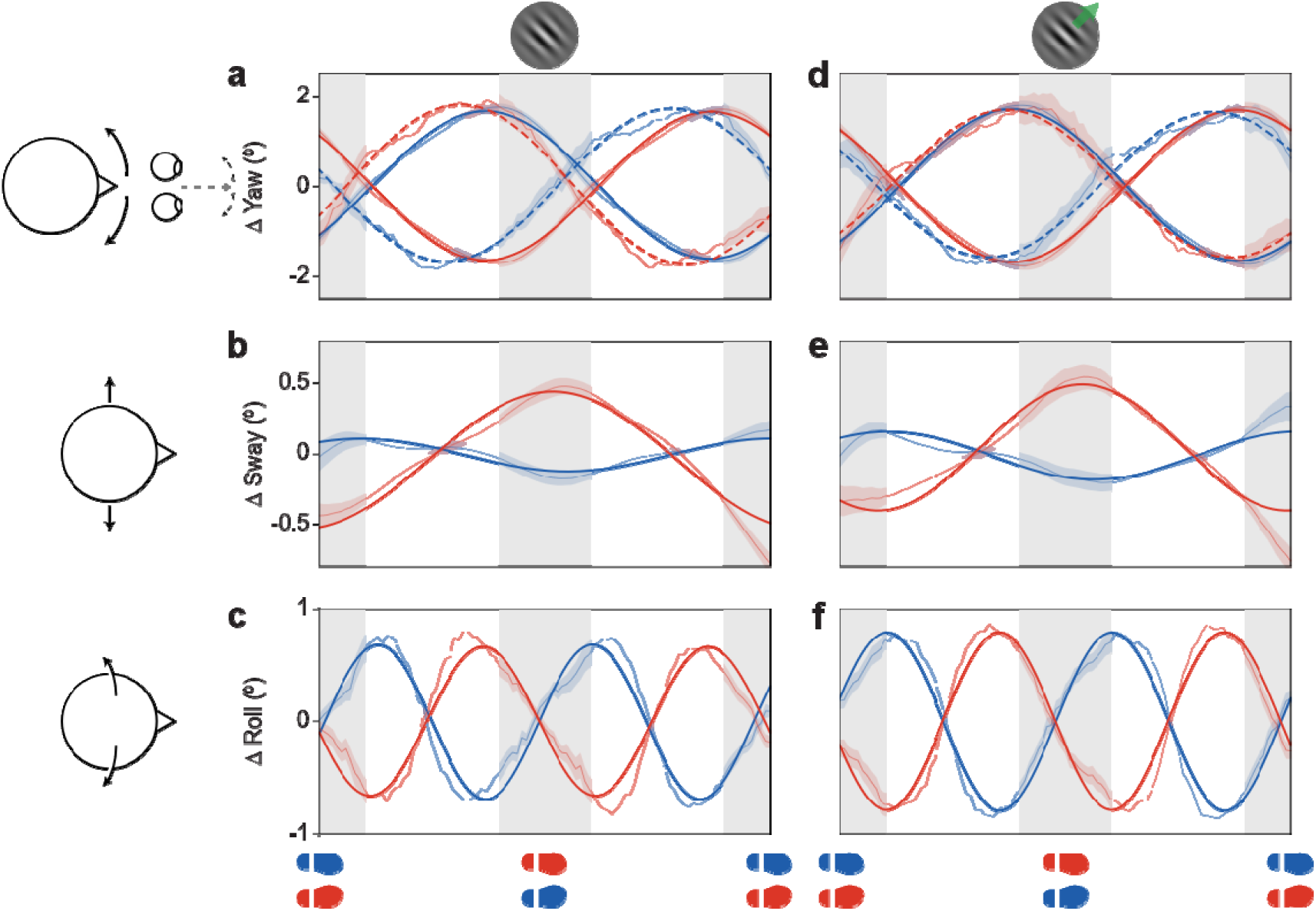
Yaw, sway, and roll across stride cycle. (**a**) Group-level (*N* = 28) change in head (solid lines) and eye (dashed lines) yaw across stride cycle for the orientation identification task. The stride cycle is comprised of two steps, beginning mid stance phase. Strides beginning with a supporting left foot are displayed in blue and those beginning with the right foot supporting are displayed in red. (**b**) Same as (**a**), but for the change in head sway only. (**c**) Same as (**b**), but for the roll in head orientation. (**d**-**f**) Same as (**a-c**), but for the group-level (*N* = 17) data from the motion direction identification task. Best fitting raised cosine functions are overlaid for (**a, b, d, e**). Shaded regions indicate ±1 SEM. Light and dark grey columns indicate estimated stance and swing phases of the stride cycle, respectively.

**Figure S3.**
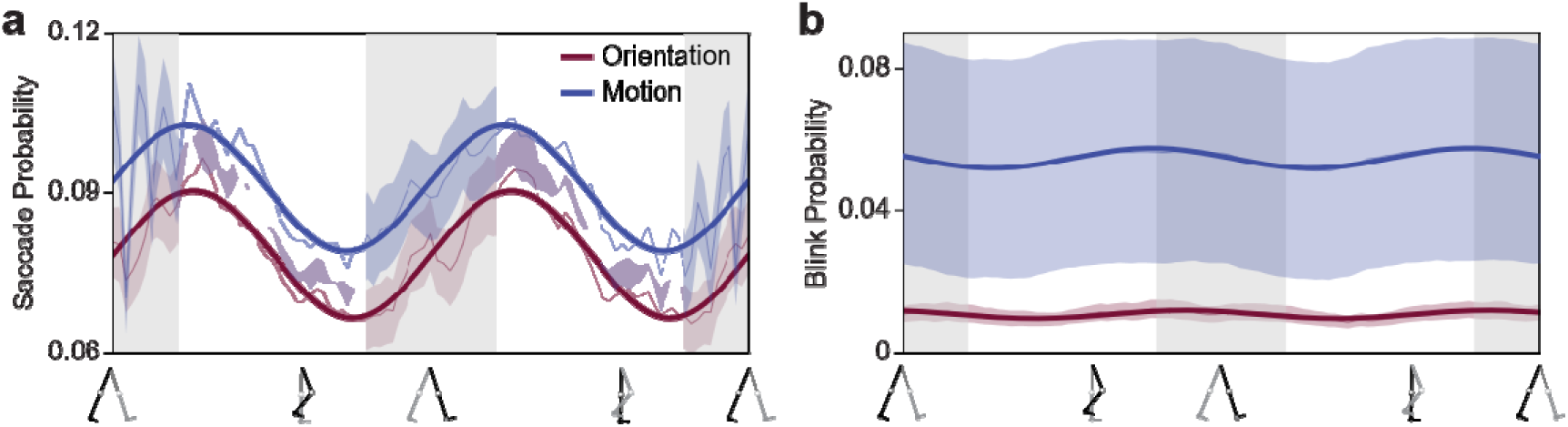
Probability of saccades and blinks across stride cycle. (**a**) Group-level distribution of saccades across the stride cycle for the orientation identification (*N* = 28) and motion direction identification tasks (*N* = 17). The stride cycle is comprised of two steps, beginning mid stance phase. (**b**) Same as (**a**), but for the distribution of blinks. Best fitting raised cosine functions are overlaid. Shaded regions indicate ±1 SEM. Light and dark grey columns indicate estimated stance and swing phases of the stride cycle, respectively.

**Figure S4.**
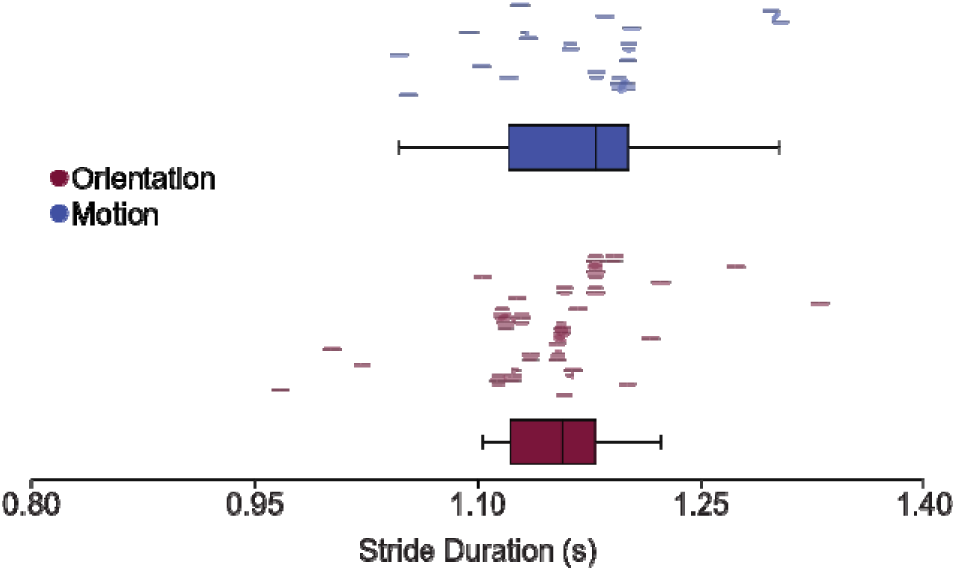
Participant stride duration. Individuals’ mean stride duration for the orientation identification and motion identification tasks. Individual error bars indicate ±1 SD. Summary boxplots are displayed below each sample.

**Figure S5.**
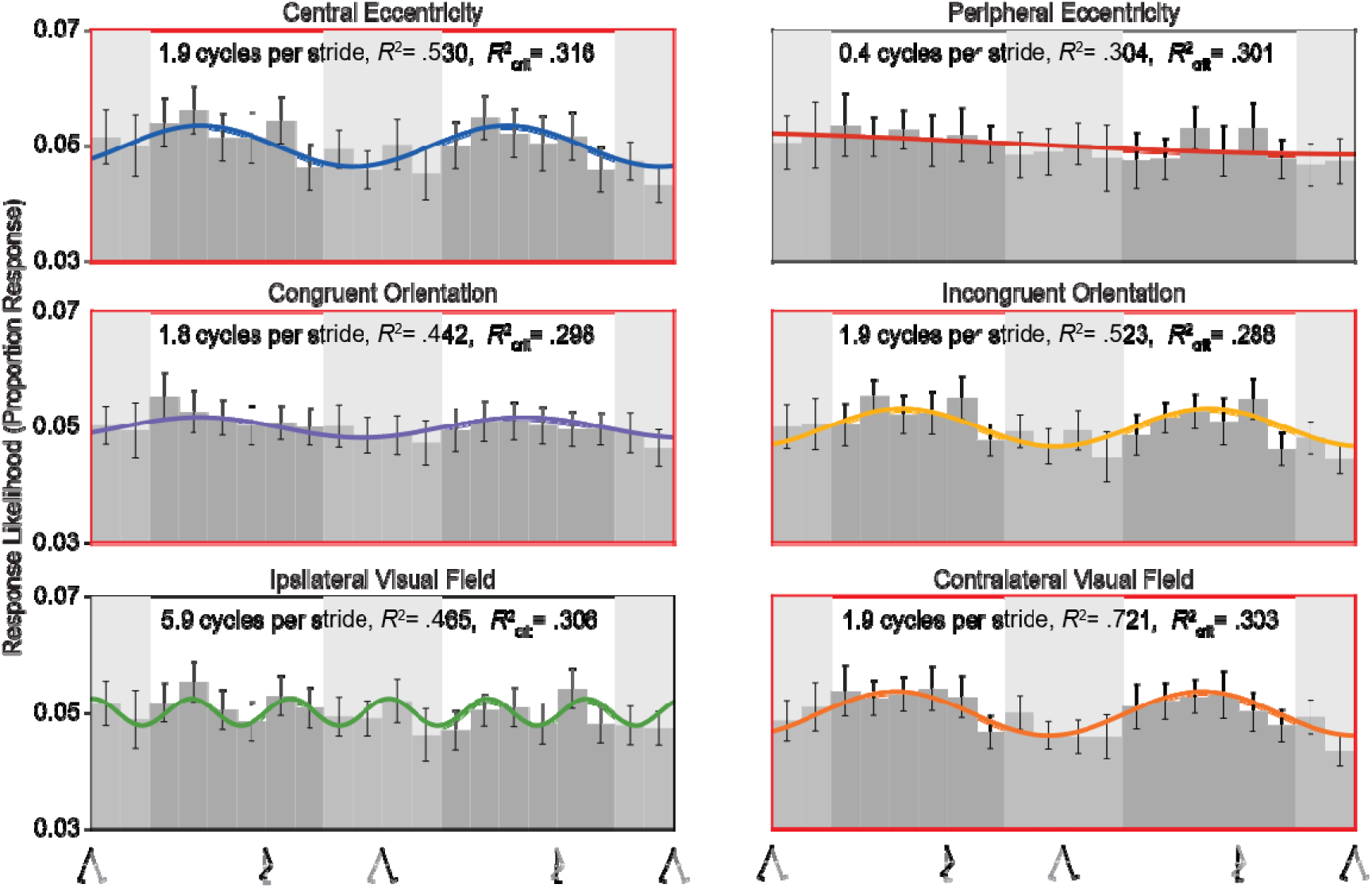
*Response likelihood for orientation identification across stride cycle*. Group-level (*N* = 28) response distribution for the orientation identification task broken down by six target conditions; centrally presented, peripherally presented, congruently oriented to the supporting foot, incongruently oriented, presented in visual hemifield ipsilateral to the supporting foot, and presented in contralateral visual hemifield. Best fitting significant raised cosine fits are overlaid. Those significant fits in the frequency range of interest (1.5 – 2.5 cycles per stride) are bordered in red. Error bars indicate ±1 SEM. Light and dark grey columns indicate estimated stance and swing phases of the stride cycle, respectively.

**Figure S6.**
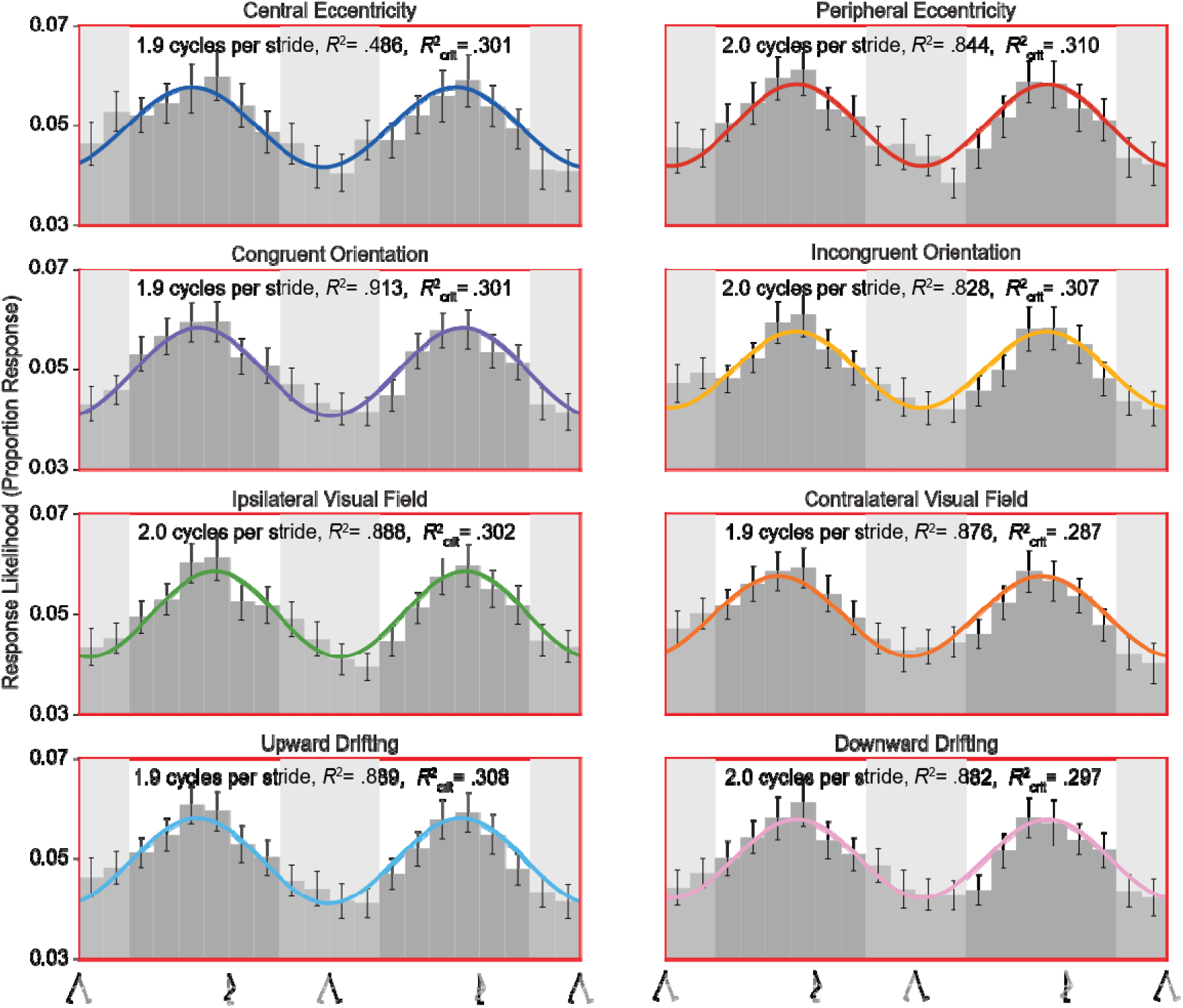
Response likelihood for motion direction identification across stride cycle. Group-level (N = 17) response distribution for the motion direction identification task broken down by eight target conditions; centrally presented, peripherally presented, congruently oriented to the supporting foot, incongruently oriented, presented in visual hemifield ipsilateral to the supporting foot, presented in contralateral hemifield, upwardly drifting, and downward drifting. Best fitting significant raised cosine fits are overlaid. Those significant fits in the frequency range of interest (1.5 – 2.5 cycles per stride) are bordered in red. Error bars indicate ±1 SEM. Light and dark grey columns indicate estimated stance and swing phases of the stride cycle, respectively.

**Figure S7.**
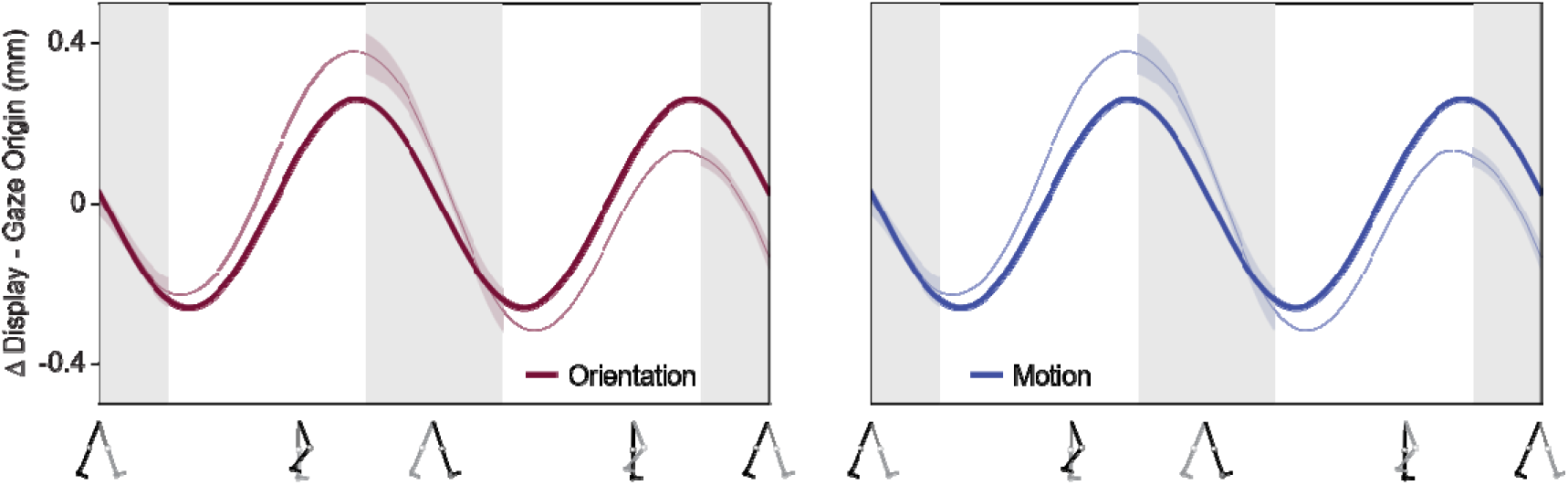
Distance from display across stride cycle. Mean distance between observers’ gaze origin and simulated display screen for the orientation identification (*N* = 28) and motion direction identification (*N* = 17) tasks. Best fitting raised cosine functions are overlaid. Shaded regions indicate ±1 SEM. Light and dark grey columns indicate estimated stance and swing phases of the stride cycle, respectively.

## References

Albright, T. D., & Stoner, G. R. (1995). Visual motion perception. Proceedings of the National Academy of Sciences, 92(7), 2433–2440. 10.1073/pnas.92.7.2433

Ambrosi, P., Burr, D. C., & Cicchini, G. M. (2022). Ideal observer analysis for continuous tracking experiments. Journal of Vision, 22(2), 3. 10.1167/jov.22.2.3

Anstis, S., Verstraten, F. A. J., & Mather, G. (1998). The motion aftereffect. Trends in Cognitive Sciences, 2(3), 111–117. 10.1016/S1364-6613(98)01142-5

Benedetto, A., Binda, P., Costagli, M., Tosetti, M., & Morrone, M. C. (2021). Predictive visuo-motor communication through neural oscillations. Current Biology, 31(15), 3401–3408.e4. 10.1016/j.cub.2021.05.026

Burr, D. C., & Wijesundra, S.-A. (1991). Orientation discrimination depends on spatial frequency. Vision Research, 31(7), 1449–1452. 10.1016/0042-6989(91)90064-C

Clark, D. J. (2015). Automaticity of walking: Functional significance, mechanisms, measurement and rehabilitation strategies. Frontiers in Human Neuroscience, 9, 246. 10.3389/fnhum.2015.00246

Clouston, G., Davidson, M. J., & Alais, D. (2025). Perception of audio-visual synchrony is modulated by walking speed and step-cycle phase. Perception, 55(5), 474–491. 10.1177/03010066251398424

Davidson, M. J. (2023). Peripersonal tracking accuracy during locomotion. 10.17605/OSF.IO/JDPWC

Davidson, M. J., Verstraten, F. A. J., & Alais, D. (2024). Walking modulates visual detection performance according to stride cycle phase. Nature Communications, 15(1), 2027. 10.1038/s41467-024-45780-4

Del Viva, M. M., & Morrone, M. C. (1998). Motion analysis by feature tracking. Vision Research, 38(22), 3633–3653. 10.1016/S0042-6989(98)00022-4

Derrington, A. M., Allen, H. A., & Delicato, L. S. (2004). Visual Mechanisms of Motion Analysis and Motion Perception. Annual Review of Psychology, 55(Volume 55, 2004), 181–205. 10.1146/annurev.psych.55.090902.141903

Durgin, F. H., Gigone, K., & Scott, R. (2005). Perception of Visual Speed While Moving. Journal of Experimental Psychology: Human Perception and Performance, 31(2), 339–353. 10.1037/0096-1523.31.2.339

Fakche, C., & Dugué, L. (2024). Perceptual Cycles Travel Across Retinotopic Space. Journal of Cognitive Neuroscience, 36(1), 200–216. 10.1162/jocn_a_02075

Fetsch, C. R., DeAngelis, G. C., & Angelaki, D. E. (2013). Bridging the gap between theories of sensory cue integration and the physiology of multisensory neurons. Nature Reviews Neuroscience, 14(6), 429–442. 10.1038/nrn3503

Foxe, J. J., & Simpson, G. V. (2002). Flow of activation from V1 to frontal cortex in humans. Experimental Brain Research, 142(1), 139–150. 10.1007/s00221-001-0906-7

Grossman, G. E., Leigh, R. J., Abel, L. A., Lanska, D. J., & Thurston, S. E. (1988). Frequency and velocity of rotational head perturbations during locomotion. Experimental Brain Research, 70(3), 470–476. 10.1007/BF00247595

Hausdorff, J. M., Purdon, P. L., Peng, C. K., Ladin, Z., Wei, J. Y., & Goldberger, A. L. (1996). Fractal dynamics of human gait: Stability of long-range correlations in stride interval fluctuations. Journal of Applied Physiology, 80(5), 1448–1457. 10.1152/jappl.1996.80.5.1448

Ho, H. T., Leung, J., Burr, D. C., Alais, D., & Morrone, M. C. (2017). Auditory Sensitivity and Decision Criteria Oscillate at Different Frequencies Separately for the Two Ears. Current Biology: CB, 27(23), 3643–3649.e3. 10.1016/j.cub.2017.10.017

Hogendoorn, H., Verstraten, F. A. J., MacDougall, H., & Alais, D. (2017). Vestibular signals of self-motion modulate global motion perception. Vision Research, 130, 22–30. 10.1016/j.visres.2016.11.002

Holliday, I. E., Anderson, S. J., & Harding, G. F. A. (1997). Magnetoencephalographic evidence for non-geniculostriate visual input to human cortical area V5. Neuropsychologia, 35(8), 1139–1146. 10.1016/S0028-3932(97)00033-X

Hubel, D. H., & Wiesel, T. N. (1962). Receptive fields, binocular interaction and functional architecture in the cat’s visual cortex. The Journal of Physiology, 160(1), 106–154. 10.1113/jphysiol.1962.sp006837

Hwang, T.-H., & Effenberg, A. O. (2021). Head Trajectory Diagrams for Gait Symmetry Analysis Using a Single Head-Worn IMU. Sensors, 21(19), Article 19. 10.3390/s21196621

Imai, T., Moore, S. T., Raphan, T., & Cohen, B. (2001). Interaction of the body, head, and eyes during walking and turning. Experimental Brain Research, 136(1), 1–18. 10.1007/s002210000533

Jain, N., Wang, A., Henderson, M. M., Lin, R., Prince, J. S., Tarr, M. J., & Wehbe, L. (2023). Selectivity for food in human ventral visual cortex. Communications Biology, 6(1), 175. 10.1038/s42003-023-04546-2

Kanwisher, N., & Yovel, G. (2006). The fusiform face area: A cortical region specialized for the perception of faces. Philosophical Transactions of the Royal Society B: Biological Sciences, 361(1476), 2109–2128. 10.1098/rstb.2006.1934

Kharb, A., Saini, V., Jain, Y., Dhiman, S., Tech, M., & Scholar. (2011). A review of gait cycle and its parameters. IJCEM Int J Comput Eng Manag, 13.

Khosla, M., Murty, N. A. R., & Kanwisher, N. (2022). A highly selective response to food in human visual cortex revealed by hypothesis-free voxel decomposition. Current Biology, 32(19), 4159–4171.e9. 10.1016/j.cub.2022.08.009

Koch, C., & Ullman, S. (1987). Shifts in Selective Visual Attention: Towards the Underlying Neural Circuitry. In L. M. Vaina (Ed.), Matters of Intelligence: Conceptual Structures in Cognitive Neuroscience (pp. 115–141). Springer Netherlands. 10.1007/978-94-009-3833-5_5

Körmendi, J., Ferentzi, E., Weiss, B., & Nagy, Z. (2021). Topography of Movement-Related Delta and Theta Brain Oscillations. Brain Topography, 34(5), 608–617. 10.1007/s10548-021-00854-0

Lamme, V. A. F., & Roelfsema, P. R. (2000). The distinct modes of vision offered by feedforward and recurrent processing. Trends in Neurosciences, 23(11), 571–579. 10.1016/S0166-2236(00)01657-X

Lappin, J. S., Tadin, D., Nyquist, J. B., & Corn, A. L. (2009). Spatial and temporal limits of motion perception across variations in speed, eccentricity, and low vision. Journal of Vision, 9(1), 30. 10.1167/9.1.30

Lewis, P., Rosén, R., Unsbo, P., & Gustafsson, J. (2011). Resolution of static and dynamic stimuli in the peripheral visual field. Vision Research, 51(16), 1829–1834. 10.1016/j.visres.2011.06.011

Manning, C., Jones, P. R., Dekker, T. M., & Pellicano, E. (2018). Psychophysics with children: Investigating the effects of attentional lapses on threshold estimates. *Attention, Perception*, & Psychophysics, 80(5), 1311–1324. 10.3758/s13414-018-1510-2

Matthis, J. S., Muller, K. S., Bonnen, K. L., & Hayhoe, M. M. (2022). Retinal optic flow during natural locomotion. PLOS Computational Biology, 18(2), e1009575. 10.1371/journal.pcbi.1009575

Matthis, J. S., Yates, J. L., & Hayhoe, M. M. (2018). Gaze and the control of foot placement when walking in natural terrain. Current Biology, 28(8), 1224–1233.

Maunsell, J. H., & Essen, D. van. (1983). The connections of the middle temporal visual area (MT) and their relationship to a cortical hierarchy in the macaque monkey. Journal of Neuroscience, 3(12), 2563–2586. 10.1523/JNEUROSCI.03-12-02563.1983

McFarland, J. M., Bondy, A. G., Saunders, R. C., Cumming, B. G., & Butts, D. A. (2015). Saccadic modulation of stimulus processing in primary visual cortex. Nature Communications, 6(1), 8110. 10.1038/ncomms9110

McKee, S. P. (1993). Psychophysics and Perception. Perception, 22(5), 505–507. 10.1068/p220505

Morton, S. M., & Bastian, A. J. (2004). Cerebellar control of balance and locomotion. *The Neuroscientist: A Review Journal Bringing Neurobiology*, Neurology and Psychiatry, 10(3), 247–259. 10.1177/1073858404263517

Muller, L., Chavane, F., Reynolds, J., & Sejnowski, T. J. (2018). Cortical travelling waves: Mechanisms and computational principles. Nature Reviews Neuroscience, 19(5), 255–268. 10.1038/nrn.2018.20

Newsome, W. T., & Pare, E. B. (1988). A selective impairment of motion perception following lesions of the middle temporal visual area (MT). Journal of Neuroscience, 8(6), 2201–2211. 10.1523/JNEUROSCI.08-06-02201.1988

Pelah, A., & Barlow, H. B. (1996). Visual illusion from running. Nature, 381(6580), 283–283. 10.1038/381283a0

Phan, C. K., Davidson, M. J., & Alais, D. (2025). Optimal phase for central and peripheral visual detection differs within the stride-cycle. *PNAS Nexus*, pgaf270. 10.1093/pnasnexus/pgaf270

Popovych, S., Rosjat, N., Toth, T. I., Wang, B. A., Liu, L., Abdollahi, R. O., Viswanathan, S., Grefkes, C., Fink, G. R., & Daun, S. (2016). Movement-related phase locking in the delta–theta frequency band. NeuroImage, 139, 439–449. 10.1016/j.neuroimage.2016.06.052

Powell, N. V., Panfili, D., Oh, Y., Matthis, J. S., & Hayhoe, M. (2026). The role of retinal flow in walking. Journal of Vision, 26(10), 6. 10.1167/jov.26.10.6

Prieto, E. A., Barnikol, U. B., Soler, E. P., Dolan, K., Hesselmann, G., Mohlberg, H., Amunts, K., Zilles, K., Niedeggen, M., & Tass, P. A. (2007). Timing of V1/V2 and V5+ activations during coherent motion of dots: An MEG study. NeuroImage, 37(4), 1384–1395. 10.1016/j.neuroimage.2007.03.080

Prokop, T., Schubert, M., & Berger, W. (1997). Visual influence on human locomotionModulation to changes in optic flow. Experimental Brain Research, 114(1), 63–70. 10.1007/PL00005624

Raiguel, S. E., Lagae, L., Gulyàs, B., & Orban, G. A. (1989). Response latencies of visual cells in macaque areas V1, V2 and V5. Brain Research, 493(1), 155–159. 10.1016/0006-8993(89)91010-X

Raiguel, S. E., Xiao, D.-K., Marcar, V. L., & Orban, G. A. (1999). Response Latency of Macaque Area MT/V5 Neurons and Its Relationship to Stimulus Parameters. Journal of Neurophysiology, 82(4), 1944–1956. 10.1152/jn.1999.82.4.1944

Rakovac, M. (2021). On Evolution and Development of Human Gait. In V. Medved (Ed.), Measurement and Analysis of Human Locomotion (pp. 39–59). Springer International Publishing. 10.1007/978-3-030-79685-3_3

Robinson, D. A. (2022). Behavior of the saccadic system: Metrics of timing and accuracy. In T. Anastasio, J. Demer, R. J. Leigh, A. Luebke, A. J. van Opstal, L. Optican, S. Ramat, & D. S. Zee (Eds.), David A. Robinson’s Modeling the Oculomotor Control System (Vol. 267, pp. 329–353). Elsevier. 10.1016/bs.pbr.2021.10.016

Sokoliuk, R., & VanRullen, R. (2016). Global and local oscillatory entrainment of visual behavior across retinotopic space. Scientific Reports, 6(1), 25132. 10.1038/srep25132

Szekely, B., & MacNeilage, P. R. (2025). Dynamic contrast sensitivity during human locomotion. Journal of Vision, 25(13), 9. 10.1167/jov.25.13.9

Takakusaki, K., Chiba, R., Nozu, T., & Okumura, T. (2016). Brainstem control of locomotion and muscle tone with special reference to the role of the mesopontine tegmentum and medullary reticulospinal systems. Journal of Neural Transmission, 123(7), 695–729. 10.1007/s00702-015-1475-4

Takamuku, S., & Gomi, H. (2021). Vision-based speedometer regulates human walking. iScience, 24(12), 103390. 10.1016/j.isci.2021.103390

Tesio, L., & Rota, V. (2019). The Motion of Body Center of Mass During Walking: A Review Oriented to Clinical Applications. Frontiers in Neurology, 10, 999. 10.3389/fneur.2019.00999

Todd, N. P. M., Govender, S., Hochstrasser, D., Keller, P. E., & Colebatch, J. G. (2025). Distinct movement related changes in EEG and ECeG power during finger and foot movement. Neuroscience Letters, 853, 138207. 10.1016/j.neulet.2025.138207

Van Kleef, J. P., Cloherty, S. L., & Ibbotson, M. R. (2010). Complex cell receptive fields: Evidence for a hierarchical mechanism. The Journal of Physiology, 588(18), 3457–3470. 10.1113/jphysiol.2010.191452

Verstraten, F. A. J. (2026). The frustration of a small n. Perception, 55(8), 723–724. 10.1177/03010066261448712

Warren, W. H., Kay, B. A., Zosh, W. D., Duchon, A. P., & Sahuc, S. (2001). Optic flow is used to control human walking. Nature Neuroscience, 4(2), 213–216. 10.1038/84054

Watson, A. B., & Pelli, D. G. (1983). Quest: A Bayesian adaptive psychometric method. Perception & Psychophysics, 33(2), 113–120. 10.3758/BF03202828

Winter, D. (1995). Human balance and posture control during standing and walking. Gait & Posture, 3(4), 193–214. 10.1016/0966-6362(96)82849-9

Winter, D. A. (1984). Kinematic and kinetic patterns in human gait: Variability and compensating effects. Human Movement Science, 3(1), 51–76. 10.1016/0167-9457(84)90005-8

Yabe, Y., & Taga, G. (2008). Treadmill locomotion captures visual perception of apparent motion. Experimental Brain Research, 191(4), 487–494. 10.1007/s00221-008-1541-3

Zaharia, A. D., Goris, R. L. T., Movshon, J. A., & Simoncelli, E. P. (2019). Compound Stimuli Reveal the Structure of Visual Motion Selectivity in Macaque MT Neurons. eNeuro, 6(6), ENEURO.0258-19.2019. 10.1523/ENEURO.0258-19.2019

Zhang, H., Morrone, M. C., & Alais, D. (2019). Behavioural oscillations in visual orientation discrimination reveal distinct modulation rates for both sensitivity and response bias. Scientific Reports, 9(1), 1115. 10.1038/s41598-018-37918-4

Zhang, X., Zhaoping, L., Zhou, T., & Fang, F. (2012). Neural Activities in V1 Create a Bottom-Up Saliency Map. Neuron, 73(1), 183–192. 10.1016/j.neuron.2011.10.035

